# Unique molecular features and cellular responses differentiate two populations of motor cortical layer 5b neurons in a preclinical model of ALS

**DOI:** 10.1101/2021.05.20.445032

**Authors:** Maria V. Moya, Rachel D. Kim, Meghana N. Rao, Bianca A. Cotto, Sarah B. Pickett, Caroline E. Sferrazza, Nathaniel Heintz, Eric F. Schmidt

## Abstract

Many neurodegenerative diseases, such as amyotrophic lateral sclerosis (ALS), lead to the selective degeneration of discrete cell types in the CNS despite the ubiquitous expression of many genes linked to disease. Therapeutic advancement depends on understanding unique cellular adaptations that underlie pathology of vulnerable cells in the context of disease-causing mutations. Here, we employ bacTRAP molecular profiling to elucidate cell type specific molecular responses of cortical upper motor neurons in a preclinical ALS model. Using two bacTRAP mouse lines that label distinct vulnerable or resilient projection neuron populations in motor cortex, we show that the regulation of oxidative phosphorylation (Oxphos) pathways is a common response in both cell types. However, differences in the baseline expression of genes involved in Oxphos and the handling of reactive oxygen species likely lead to the selective degeneration of the vulnerable cells. These results provide a framework to identify cell type-specific processes in neurodegenerative disease.

## INTRODUCTION

Amyotrophic lateral sclerosis (ALS) is a fatal late-onset neurodegenerative disease that targets the motor pathways of the central nervous system, and for which there is no cure. After symptom onset, the disease advances rapidly with most patients experiencing progressive paralysis and eventual death within 2-5 years of diagnosis. While most cases of ALS are sporadic, about 10% of cases are inherited (Rowland and Shneider, 2001; Tandan and Bradley, 1985). Gene linkage, GWAS, and high throughput sequencing studies have identified disease-causing mutations in over 50 genes, including *SOD1*, *C9ORF72*, *TARDBP, FUS, OPTN, and TBK1* (DeJesus-Hernandez et al., 2011; Kwiatkowski et al., 2009; Renton et al., 2011; Rosen et al., 1993; Sreedharan et al., 2008; Vance et al., 2009). Most ALS-linked genes are expressed widely throughout the brain and periphery and are involved in a variety of cellular pathways including RNA processing, autophagy, membrane trafficking, and antioxidant function (Cook and Petrucelli, 2019). Despite their ubiquitous expression, mutations in these genes lead to specific loss of discrete cell populations, including the lower motor neurons in the brainstem and spinal cord, and upper motor neurons in the motor cortex. This selective loss brings to question which properties of motor neurons make them specifically vulnerable to ALS-causing mutations while other cells are seemingly resistant. Determining the key cell type specific biological contributors to cellular pathology and vulnerability will thereby lay the groundwork for the discovery of more efficacious therapies.

Pathology and dysfunction of both lower motor neuron and upper motor neurons is required for a diagnosis of ALS (Braak et al., 2013; Ravits et al., 2007). However, the majority of preclinical research into understanding motor neuron-specific pathology in ALS has focused on lower motor neurons (see Ragagnin et al., 2019). This is because they are anatomically segregated from other cell types and have distinctive morphological features and molecular markers, facilitating their identification and targeting (Borges and Iversen, 1986; Ichikawa et al., 1997), and can be differentiated and grown in vitro (Sances et al., 2016). Accessing the vulnerable cells in the forebrain and cortex has been more challenging due in large part to the heterogeneity of cortical neuron types and a relative lack of reliable markers to distinguish between them. Postmortem brains from ALS patients show a loss of spinal-projecting Betz cells in layer 5 (L5) of primary motor cortex (M1; Hammer et al., 1979) that is accompanied by vacuolization of apical dendrites (Genc et al., 2017; Saberi et al., 2015), and hyperexcitability of cortical neurons and their axons (Eisen et al., 1996; Kohara et al., 1996; Vucic et al., 2013; Zanette et al., 2002). In preclinical rodent models, L5 cells in M1, in particular layer 5b (L5b) corticospinal neurons, show early hyperexcitability along with changes in dendritic structure and spine density (Fogarty et al., 2015; Kim et al., 2017; Saba et al., 2015). These are accompanied by corticospinal tract degeneration (Thomsen et al., 2014), dendritic vacuolization, and eventually death (Ozdinler et al., 2011; Yasvoina et al., 2013; Zang and Cheema, 2002). Many of these studies relied on acute labeling methods, such as retrograde tracing, or transgenic animals that broadly labeled excitatory L5b neurons. However, recent descriptions of axonal projection diversity and molecular heterogeneity within L5b necessitate that ALS pathology in these deep layer neurons be assessed reproducibly with even greater cell type resolution.

Within the motor areas of cortex, multiple subpopulations of L5b pyramidal tract-projecting (PT) neurons have been distinguished by their closely related but unique molecular profiles and axonal projection targets (Economo et al., 2016; Gerfen et al., 2018; Oswald et al., 2013; Tasic et al., 2016; Tasic et al., 2018). Previously, these L5b PT cell types were very difficult to distinguish from each other-in part because of their close laminar proximity and similar morphological features- and were therefore challenging to dissociate molecularly and anatomically in ALS mouse models. However, understanding the biological differences between these closely related PT neuron types can provide a unique opportunity to describe the specific features that may confer L5b neuron vulnerability in ALS and contribute to their pathology during disease progression.

Here, we utilized bacTRAP transgenic mice to target distinct populations of PT neurons in L5b of M1, including a subset that exhibits vulnerability to degeneration in the SOD1*G93A mouse model of ALS (Chiu et al., 1995; Gurney et al., 1994). We employed Translating Ribosome Affinity Purification (TRAP; Heiman et al., 2014; Heiman et al., 2008) to determine molecular features that distinguish these L5b cell types and may confer differential vulnerability in disease. We show that only the Gprin3-TRAP-labelled corticospinal-projecting cells located in lower L5b (LL5b) degenerate in the SOD1*G93A mice while upper layer 5b (UL5b) pons-projection cells labeled in Colgalt2-TRAP mice are spared. Gprin3 cells exhibited a more robust molecular response to SOD1*G93A expression, including modulation of genes associated with maintenance of axon and synapse structure and function, while both resistant Colgalt2-TRAP cells and vulnerable Gprin3-TRAP cells increase their expression of oxidative phosphorylation (Oxphos) genes. Lastly, we reveal that Gprin3 cells show baseline differences in expression of oxidative stress and antioxidant response transcription factors, and that these factors and their targets were further changed in SOD1*G93A mice.

## METHODS

### Animals

All animal procedures and experiments were done with approval from The Rockefeller University Institutional Animal Care and Use Committee (IACUC) and in accordance with National Institutes of Health (NIH) policies and guidelines.

Colgalt2-bacTRAP DU9 and Gprin3-bacTRAP ES152 mice, as well as TRAP lines for other cortical cell types, were generated using the two plasmid/one recombination method (Gong et al., 2010) at The Rockefeller University. Generation of Colgalt2-bacTRAP mice were described previously (Doyle et al., 2008). A sequence homology arm corresponding to the region upstream of the ATG start codon of *Gprin3* was cloned into the pS296 targeting vector containing EGFP-L10a (Heiman et al., 2008). Recombination was performed by electroporating the modified pS296 vector into competent DH10β bacteria containing a pSV1.RecA plasmid and the RP24-127P5 BAC. The modified BAC was isolated and microinjected into the pronuclei of fertilized FVB/N mouse oocytes at 0.5 ng/μl. Transgenic founder mice were generated and crossed to C57BL/6J mice. F1 progeny were screened for proper transgene expression by EGFP genotyping and immunohistochemistry. Expression in L5b was used as a benchmark for transgenic line expansion. C57Bl/6J (Jackson Laboratory, #000664) were used for breeding in this study. The Tg(SOD1*G93A)1Gur mice (Jackson Laboratory, #002726) were used for ALS experiments.

### Rotarod

SOD1*G93A mice and wildtype littermates (WT) were trained on the apparatus for three days beginning at five weeks of age (about postnatal day 35, ~P35) and then tested once weekly for six weeks. During the test, acceleration was increased from 5-18 rpm over the course of 180 seconds and the latency to fall was measured for two trials. Falls were detected automatically by a sensor at the base of the apparatus.

### CTB injections

Animals were anesthetized via IP injection of 1% Ketamine/0.1% Xylazine (doses 1 mL/kg and 0.1 mL/kg respectively) in 0.9% saline. Following anesthesia, animals were positioned on the stereotaxic apparatus, and skin and periosteum were incised above the skull. Pons injections were made at AP −4.1 mm, ML +/−0.6 mm, DV −5.2 mm. Holes were drilled in the skull at the target coordinates using a dental drill. Cholera toxin beta (Alexa 555-conjugated CTB, Thermo Fisher) was injected using a Hamilton syringe with a pulled glass capillary pipette at a rate of 0.1 μL/min to a total volume of 0.25 μL. Incisions were closed using Vetbond (3M). For spinal cord injections, an incision was made from the base of the neck down between the shoulder blades to target the cervical vertebrae. Back and shoulder muscles were separated by cutting connective ligaments with a scalpel. Once the spinal column was exposed, connective tissue was pushed aside slowly using the scalpel blade. Using forceps and surgical scissors, the C6 vertebrae was cut away to reveal the dura and spinal tissue. After carefully cutting a hole in the dura using forceps and scalpel, CTB was injected at a rate of 0.1 μL/min to a total of 0.15-0.20 μL at the following coordinates: AP C6 vertebra, ML +/−0.4 mm, DV −1.1 mm. Sutures were used to reposition back and shoulder muscles before suturing the incision. Animals were allowed to recover in clean cages over a warming plate and were given low dose ibuprofen in drinking water during recovery. Animals were perfused after at least 48 hrs following surgery to allow tracer to fully travel the length of the axons.

### Tissue collection and histology

Brain and spinal cord tissue were collected from experimental mice following cardiac perfusion. Mice received an IP injection of 1% Ketamine/0.1%Xylazine (lethal doses 2 mL/kg and 0.2mL/kg respectively). Mice were perfused first with approximately 40 mL of sterile phosphate buffered saline (PBS) pH 7.4, followed by approximately 30mL of freshly made 4% paraformaldehyde in PBS. Brain and spinal cord were removed and placed in 4% paraformaldehyde overnight at 4°C with shaking. Tissues were then transferred to 30% w/v sucrose in PBS and stored at 4°C for approximately 2 days. Brain and spinal cord tissues were sectioned on a sliding freezing microtome at a thickness of 40μm. To section spinal cord tissue on the microtome, tissue was first embedded in Neg 50 (Thermo Fisher) or OCT medium (Tissue-Tek). Sections were stored in non-freezing storage media (25% w/v ethylene glycol, 25% glycerol, 50% PBS) at −20°C until further processing. Before staining or mounting, sections were washed 3 × 5 min with PBS.

### Immunostaining

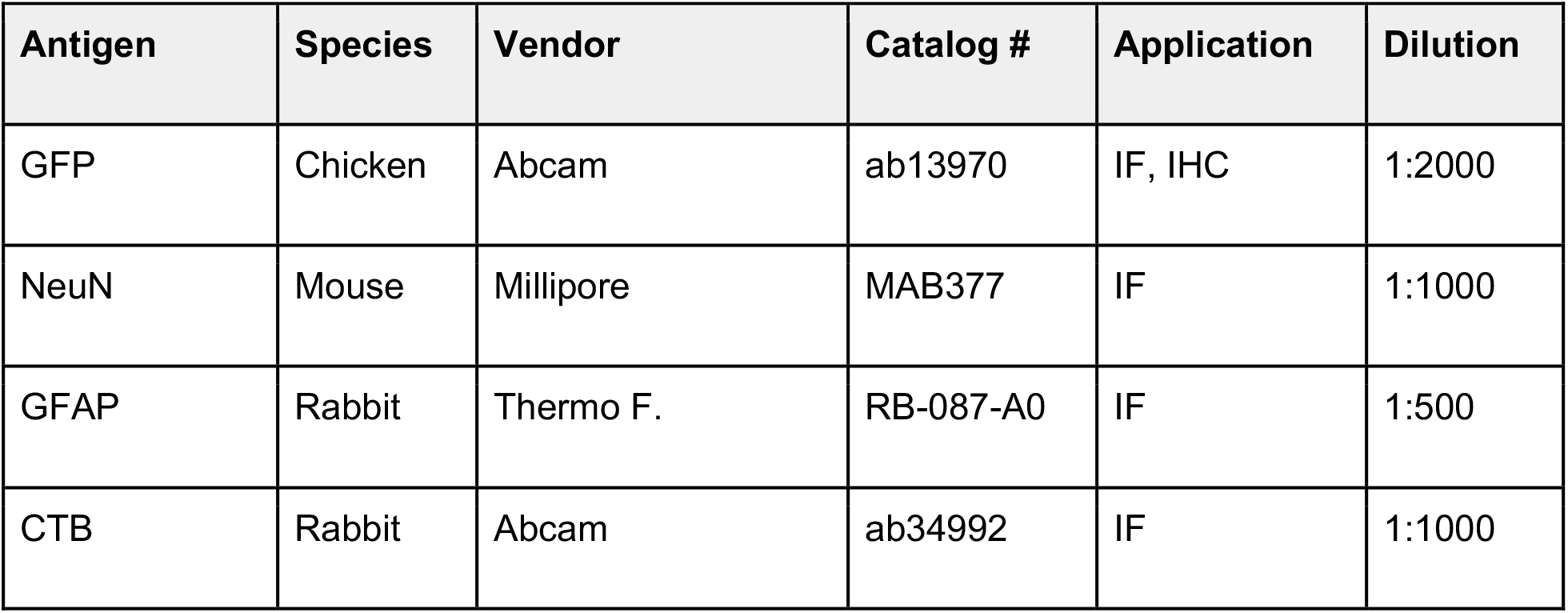
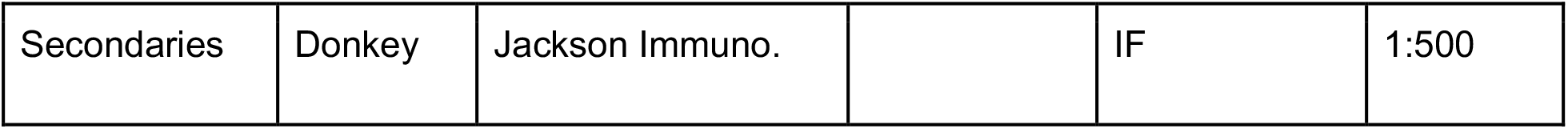
The following antibodies were used in study:

| Antigen | Species | Vendor | Catalog # | Application | Dilution |
| --- | --- | --- | --- | --- | --- |
| GFP | Chicken | Abcam | ab13970 | IF, IHC | 1:2000 |
| NeuN | Mouse | Millipore | MAB377 | IF | 1:1000 |
| GFAP | Rabbit | Thermo F. | RB-087-A0 | IF | 1:500 |
| CTB | Rabbit | Abcam | ab34992 | IF | 1:1000 |
| Secondaries | Donkey | Jackson Immuno. |  | IF | 1:500 |

All fluorescent antibody staining experiments were performed on 40μm free-floating tissue sections as follows: sections were washed in PBS pH 7.4, 3 × 5 min to remove anti-freezing storage solution. They were then incubated with 2.5% Normal Donkey serum (NDS, Jackson Immuno) with 2.5% TritonX-100 in PBS for approximately 30 mins, followed by 1% NDS, 1% TritonX-100, and primary antibodies in PBS overnight at room temperature with shaking. Sections were washed with PBS 3 × 5 min before incubating in 1% NDS with fluorescent secondary antibodies in PBS for approximately 1 hour. Sections were washed with PBS 3 × 5 min before being mounted on slides and coverslipped with Prolong Gold with DAPI (Thermo). Sections were then visualized using a confocal microscope.

For non-fluorescent immunohistochemistry with DAB, a 10 min incubation in 0.3% hydrogen peroxide in PBS and 3 × 5 min PBS washes preceded the blocking/permeabilization step. A biotinylated secondary antibody was used, followed by a 1 hr incubation in ABC (Vector Labs). DAB (Vector Labs) was then used to deposit the stain at the site of antigen. Sections were then visualized using a brightfield microscope.

### RNAScope in situ hybridization (ISH)

Cryostat sections (14 μm) for FISH experiments were made from PFA perfusion-fixed and 30% sucrose cryopreserved brains that were embedded in OCT freezing media. On-slide labeling was performed using RNAscope ISH technology and its associated kits. Probes against *Vat1l*, *Lypd1*, and *Nefh* (ACD Biotech, #495401, #318361, #443671) were used to label L5b cells in Colgalt2 and Gprin3 sections using the RNAscope Multiplex Fluorescent Reagent Kit v2 (ACD Biotech, #323100). All steps were carried out according to manufacturer recommendations in protocol #323100-USM, available on the manufacturer website.

Immunostaining after ISH was performed on-slide as described above, and concentrations for serums, Triton, and antibodies, as well as incubation times, were identical to free-floating immunostaining protocol.

### Microscopy

A Zeiss Axio Observer.Z1 with LSM 700 confocal microscope equipped with 50 mW, 400-640 nm lasers was used for all fluorescent visualizations. Zen software (Zeiss) was used for acquisition of images. FIJI (ImageJ, SciJava) was used for brightness and contrast adjustments, as well as image analysis (Rueden et al., 2017; Schindelin et al., 2012; Schneider et al., 2012). Tiled, brightfield images of whole DAB-stained sections were acquired using a Zeiss Axio Imager.M2 and Neurolucida (Micro Bright Field).

### Image analyses

#### Multi-channel signal co-localization

For co-localization analyses, position markers were placed in FIJI over any cells that appeared subjectively positive for a marker. The position markers were saved as ROIs in the ROI manager and placed over the next channel image of interest. The number of points that were positive in this second channel was counted as being double-positive. This analysis framework was used for CTB/GFP and PRV-mCherry/GFP co-localization in retrograde tracing, and for L5b marker co-localization in mouse and human tissues.

#### Cell counts

For determining loss of GFP+ cells in SOD1*G93A mice, a 3 × 2 × 3 (w × h × z) z-stack tile scan of 20X fields was acquired on the confocal microscope. Every 6th section between +1.00 mm and +0.00 mm to Bregma was imaged (one section every ~240 μm), with each hemisphere being imaged separately. The z-stacks were collapsed in FIJI using the maximum intensity projection. Cells were then counted only if apical dendrite was clearly visible. All counting was performed blind to disease condition. Data are presented as percent of WT mean at each Bregma coordinate along the AP extent of M1.

#### Cell depth analyses

Cell depths were determined using a custom script (github.com/mvmoya) written for FIJI/ImageJ that calculates the percentage depth from pia and white matter. The pial and white matter surfaces were manually delineated, and the points were placed on the image to mark the locations of the cells. The algorithm then drew a reference line perpendicular to the pial line that intersected the cell’s point and terminated at the intersection with the white matter line. The cell’s percent distance from the pial line along the reference line was then calculated and reported as a percentage depth from pia.

### TRAP and RNA-Seq analyses

#### TRAP

TRAP experiments from cortical cell types were performed as previously described (Heiman et al., 2014; Nectow et al., 2017), with minor adaptations. For cell type characterizations, ~P40 animals were sacrificed, and M1 cortex sub-dissected in ice-chilled HBSS containing 2.5 mM HEPES-KOH (pH 7.4), 35 mM glucose, 4 mM NaHCO3, and 100 μg/ml cycloheximide. Cortices from 3 animals were pooled per biological replicate, with 3 replicates per cell type being prepared in total (9 animals). Samples were homogenized in extraction buffer containing 10 mM HEPES-KOH (pH 7.4), 150 mM KCl, 5 mM MgCl2, 0.5 mM DTT, 100 μg/ml cycloheximide, RNasin (Promega) and SUPERase-In (Life Technologies) RNase inhibitors, and Complete-EDTA-free protease inhibitors (Roche), and then centrifuged at 2000 × g to clear lysate debris. IGEPAL CA- 630 (NP-40, Sigma) and DHPC (Avanti Polar Lipids, Alabaster, AL) were added to the resulting supernatant to a concentration of 1% each, followed by 20,000 × g centrifugation. Polysomes were immunoprecipitated (IP’ed) using 100 μg custom monoclonal anti-GFP antibodies (50 μg of clone 19C8 with 50 μg of clone 19F7) bound to biotinylated Protein L (Pierce, Thermo Fisher) coated streptavidin-coated magnetic beads (Thermo Fisher), and washed with high salt buffer containing 10 mM HEPES-KOH (pH 7.4), 350 mM KCl, 5 mM MgCl2, 1% IGEPAL CA- 630, 0.5 mM DTT, 100 μg/ml cycloheximide, and RNasin RNase inhibitors (Promega). Overnight IP’s were eluted and purified using Absolutely RNA Nanoprep kit (Agilent). RNA quality was assessed using Nanodrop spectrophotometer and Agilent 2100 Bioanalyzer. Samples with RNA integrity values > 7 were used to prepare libraries for sequencing. RNAs from TRAP IPs (15 ng per sample) were converted to cDNA using the Nugen Ovation RNA-Seq System V2 kit. cDNAs were then fractionated by sonication and libraries were made using the Illumina TrueSeq RNA Sample Preparation kit v2, following manufacturer’s instructions. Sequencing was performed at The Rockefeller University Genomics core using the Illumina HiSeq 2500 platform (Illumina). For disease TRAP cohorts, whole cortex (Colgalt2 samples) or M1 cortex (M1 input and Gprin3 samples) from the ~P110 (symptomatic) timepoint were collected from SOD1*G93A-crossed Colgalt2- and Gprin3-bacTRAP animals and healthy littermates, with each individual used as a separate replicate. Cortices were then homogenized, and polysomes were IP’ed as described above. All other library preparation and sequencing procedures were also performed as described above, with the exception of our Colgalt2 ALS cohort, which was sent to the NYGC for sequencing.

#### Sequencing analyses

Sequence and transcript coordinates for mouse mm10 UCSC genome and gene models were retrieved from the Bioconductor Bsgenome.Mmusculus.UCSC.mm10 (version 1.4.0) and TxDb.Mmusculus.UCSC.mm10.knownGene (version 3.4.0) Bioconductor libraries respectively. RNA-seq reads are aligned to the genome using the subjunc method (version 1.30.6) in Rsubread (Liao et al., 2013) and exported as bigWigs normalized to reads per million using the rtracklayer package (version 1.40.6). Counts in genes were generated using featureCounts within the Rsubread package (version 1.30.6) with default settings and the TxDb.Mmusculus.UCSC.mm10.knownGene gene models. Differential expression analysis on whole gene counts was performed using all default parameters in DESeq2 version 1.26.0 in RStudio version 1.2.5033 (Love et al., 2014) http://www.rstudio.com/, with R version 3.6.3. Gene ontology enrichment analyses were performed using Metascape (Zhou et al., 2019) with default parameters, with genes with mean counts per million > 100 being used for these analyses. GO categories used for functional category expression analyses included GO:1990542 mitochondrial transmembrane transport, GO:0006979 response to oxidative stress, GO:0016209 antioxidant activity, GO:0006119 oxidative phosphorylation, and GO:0006099 tricarboxylic cycle.

## RESULTS

### Colgalt2-TRAP and Gprin3-TRAP cells occupy distinct sublayers of L5b in the primary motor cortex

We first set out to generate transgenic mouse lines to target PT neurons in L5b of the primary motor cortex (M1). The GENSAT database (Gong et al., 2003) and the Allen Brain Atlas (Lein et al., 2007) were used to generate a list of candidate genes that showed selective expression in L5b pyramidal populations. Bacterial artificial chromosomes (BACs) were then used to generate transgenic bacTRAP mice expressing the EGFP-tagged ribosomal protein L10a (EGFP-L10a) under the control of these cell type-specific promoters. Two of the candidate genes, *Colgalt2* and *Gprin3*, drove EGFP-L10a expression in cortical pyramidal neurons found across different regions of cortex, including the premotor and motor areas (**Figure S1A-D**). In the Colgalt2-TRAP line EGFP-L10a expression was apparent across a broad medial-lateral (ML) and anterior-posterior (AP) extent across all cortical regions (Doyle et al., 2008; Groh et al., 2010; Schmidt et al., 2012). EGFP-L10a expression in the Gprin3-TRAP line was also observed in a small subset of small superficial layer interneurons, though more GFP+ L5b pyramidal neurons were present primarily in rostromedial cortical areas, with few GFP+ pyramidal neurons detected posterior to Bregma, lateral to sensory cortical areas, and medial to secondary motor cortex. Colgalt2 cells were rarely observed in subcortical regions. In contrast, Gprin3 cells were also detected in the hippocampus, the striatum, and other subcortical areas.

To confirm localization of Colgalt2-TRAP and Gprin3-TRAP cells to L5b, the depth of EGFP+ cells was measured relative to the pial surface (**Figures 1A, B** and **S1E, F**). In M1, Colgalt2 cells were found in a narrow ~200 μm layer between ~40-65% depth of the cortex. They were located slightly deeper than S100a10-TRAP cells in L5a, suggesting that Colgalt2 cells were located in an UL5b. Gprin3 cells were instead found at ~55-75% depth and were located superficial to Ntsr1-TRAP cells in L6a, suggesting that Gprin3 cells reside in the LL5b (**Figure 1B**). In primary sensory cortex (S1), Colgalt2 and Gprin3 cells were found at the same depth, ~50-70% from the pial surface, indicating that the division of Colgalt2-TRAP and Gprin3-TRAP neurons into upper and lower sublayers is unique to motor areas. Both Colgalt2 and Gprin3 neurons displayed larger cell body size compared to other deep layer neurons, including L5a S100a10 and L6a Ntsr1 cells (**Figure 1C**), as well as to other randomly selected pyramidal neurons across all cortical layers (**Figure S1G**). Immunofluorescent staining of tissue from each bacTRAP line revealed 79% of Colgalt2 and 94% of Gprin3 EGFP+ cells were co-labeled with Ctip2, a known marker of subcortical-projection neurons, confirming the PT identity of both populations (**Figures 1D** and **S1H**).

**Figure 1.**
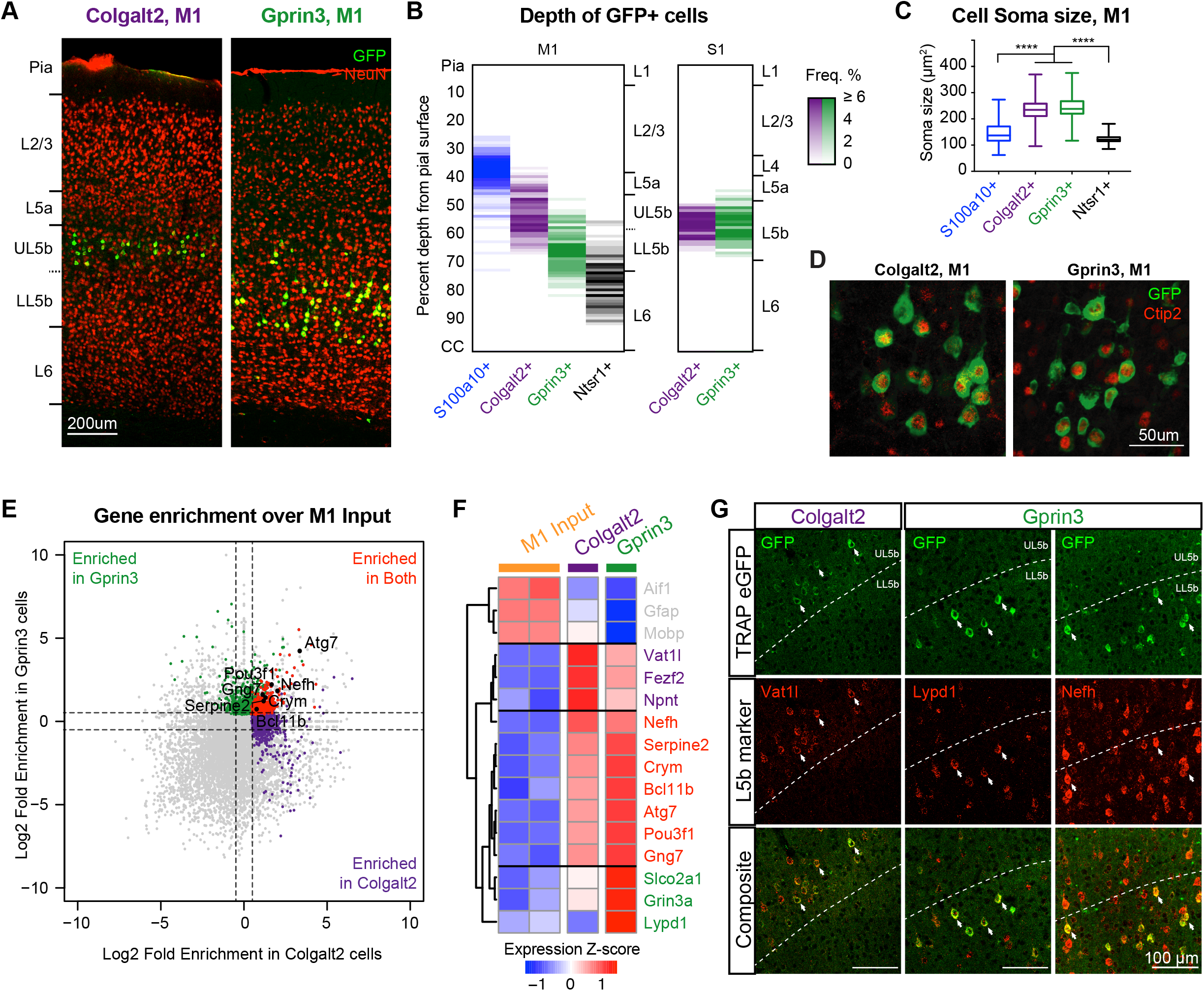
Colgalt2 and Gprin3 bacTRAP lines label molecularly and anatomically distinct populations projection neurons in layer 5b of M1. **A.** Anti-GFP immunostaining (green) showing expression of eGFP-L10a in Colgalt2-bacTRAP DU9 (left) and Gprin3-bacTRAP ES152 (right) animals present in deep layer cells of primary motor cortex (M1). Red staining indicates NeuN+ cells. Scale bar = 200 μm. **B.** Frequency distribution of laminar depth of labeled cells in Colgalt2-bacTRAP (purple, n = 209 cells), Gprin3-bacTRAP (green, n = 163 cells), S100a10-bacTRAP (blue, n = 248 cells) and Ntsr1-bacTRAP (black, n = 146 cells) mice in M1 and in neighboring S1 (L5b populations only, Colgalt2-bacTRAP, n = 125; Gprin3-bacTRAP, n = 95 cells). **C.** Box and whisker plots of population distribution for soma sizes (area in μm^2^) of S100a10 (blue, mean ± SEM 144.2 ± 1.7 μm^2^), Colgalt2 (purple, 233.3 ± 2.6 μm^2^), Gprin3 (green, 244.0 ± 2.7 μm^2^), and Ntsr1 (black, 122.3 ± 1.3 μm^2^) cells. **** p < 0.0001 by one-way ANOVA and subsequent Tukey multiple comparison test. **D.** Immunostaining for Ctip2 (red) and GFP (green) in Colgalt2-bacTRAP (left) and Gprin3-bacTRAP (right) animals shows overlap of expression. See also Figure S1H. **E.** Scatterplot showing Log_2_ fold enrichment values from DE analyses of Colgalt2 TRAP vs. M1 input (x-axis) and Gprin3 TRAP vs. M1 input (y-axis). Genes enriched in Colgalt2 only (purple), Gprin3 only (green), or both (red) are indicated. L5b PT cell marker genes (black) are labeled. **F.** Heatmap showing relative expression of glial genes (gray), and a subset of known L5b marker genes that showed enrichment in either Colgalt2 cells only (purple), both cell types (red), or Gprin3 cells only (green) across M1 input, Colgalt2 TRAP, and Gprin3 TRAP data sets. Values reported as z-scores of normalized CPM, averaged across biological replicates, and scaled for each gene. **G.** Images showing FISH for *Vat1l* in UL5b (left column), *Lypd1* in LL5b (middle column), and *Nefh* in both sublayers (right column) co-labeled with anti-GFP immunofluorescence in M1 of bacTRAP mice. Arrows indicate cells that were double-positive for GFP and the probed marker gene. Scale bar = 100 μm.

We next wanted to determine whether these anatomical findings corresponded to underlying gene expression patterns of these two cell types. We employed TRAP (Heiman et al., 2014; Heiman et al., 2008) to examine the molecular profiles of each cell population more closely. EGFP-tagged polysomes containing the EGFP-L10a transgene were IP’ed from microdissected M1 homogenates from each bacTRAP line using anti-EGFP antibodies. Polysome-bound mRNA molecules were then isolated and subsequently processed for high-throughput RNA-Seq. Relative to other cell types in the cortex and whole cortex input samples, UL5b Colgalt2 and LL5b Gprin3 cells clustered closely together by principal component analysis (PCA; **Figure S2A**), suggesting that while distinct, these two cell types likely share many molecular features. To determine the genes that were enriched across both Colgalt2 and Gprin3 cells, we performed differential expression (DE) analysis comparing TRAP mRNA from each cell type to mRNA purified from whole M1 (M1 input; **Figure S2B**). A total of 1598 genes were significantly enriched (padj < 0.05 and log_2_ fold enrichment; LFE > 0.5) in M1 Colgalt2 IPs and 1568 genes were significantly enriched in Gprin3 IPs (**Figure S2B**). We found that 486 enriched genes were shared by both cell types, including known L5b marker genes, such as *Fezf2*, *Crym*, *Nefh*, and *Serpine2*, while glial markers such as *Aif1*, *Gfap*, and *Mobp* were depleted (**Figure 1E, F**). Identification of genes specifically enriched within each cell type can serve as novel markers to distinguish these two PT cell types from each other and from other cell types in motor cortex without transgenic labeling. For example, *Vat1l* displayed high levels of enrichment in Colgalt2 cells, while *Lypd1* showed higher expression in Gprin3 cells (**Figure 1F**). Indeed, in situ hybridization (ISH) revealed that *Vat1l* transcripts specifically labeled GFP+ UL5b Colgalt2 cells, *Lypd1* was expressed specifically in GFP+ LL5b Gprin3 cells, and the pan-L5b marker, *Nefh,* labeled cells across both sublayers (**Figure 1G**). Together, these data show that Colgalt2-TRAP and Gprin3-TRAP neurons represent closely related but molecularly distinct populations of PT cells located across discrete sublayers of L5b.

### Colgalt2 and Gprin3 cells share a PT projection to the pons, but Gprin3 cells alone project to more distal PT targets

Ctip2-positive pyramidal neurons located in L5b of M1 project to various subcortical areas and send axons to long-range PT targets, such as the spinal cord (Arlotta et al., 2005). We therefore asked whether M1 Colgalt2 and Gprin3 cells projected to PT targets. We injected the retrograde tracer cholera toxin β subunit (CTB) into the pons, a proximal PT target (**Figure 2A**), and C6 cervical spinal cord, a distal target (**Figure 2C**), to label neurons that project to these two structures in adult Colgalt2 and Gprin3 bacTRAP mice. In pons-injected animals, CTB+ neurons were clearly localized to both UL5b and LL5b of M1 (**Figure 2B**) with laminar distributions 40-75% from pia, overlapping with both Colgalt2 and Gprin3 cells (**Figure 2E**). In Colgalt2 bacTRAP mice, 64% of CTB+ cells in UL5b (and 38% of all CTB+ cells) were EGFP+ (**Figure 2F**), suggesting that Colgalt2 cells represent the majority of pons-projecting cells in UL5b. In Gprin3 bacTRAP animals, 77% of pontine CTB+ cells in LL5b (and 48% of all CTB+ cells) were co-labeled with EGFP, indicating that Gprin3 neurons represent the majority of LL5b cells that project to the pons (**Figure 2F**). These data also indicate that Colgalt2-TRAP and Gprin3-TRAP neurons represent the vast majority (~85%) of pons-projecting cells in M1 L5b.

**Figure 2.**
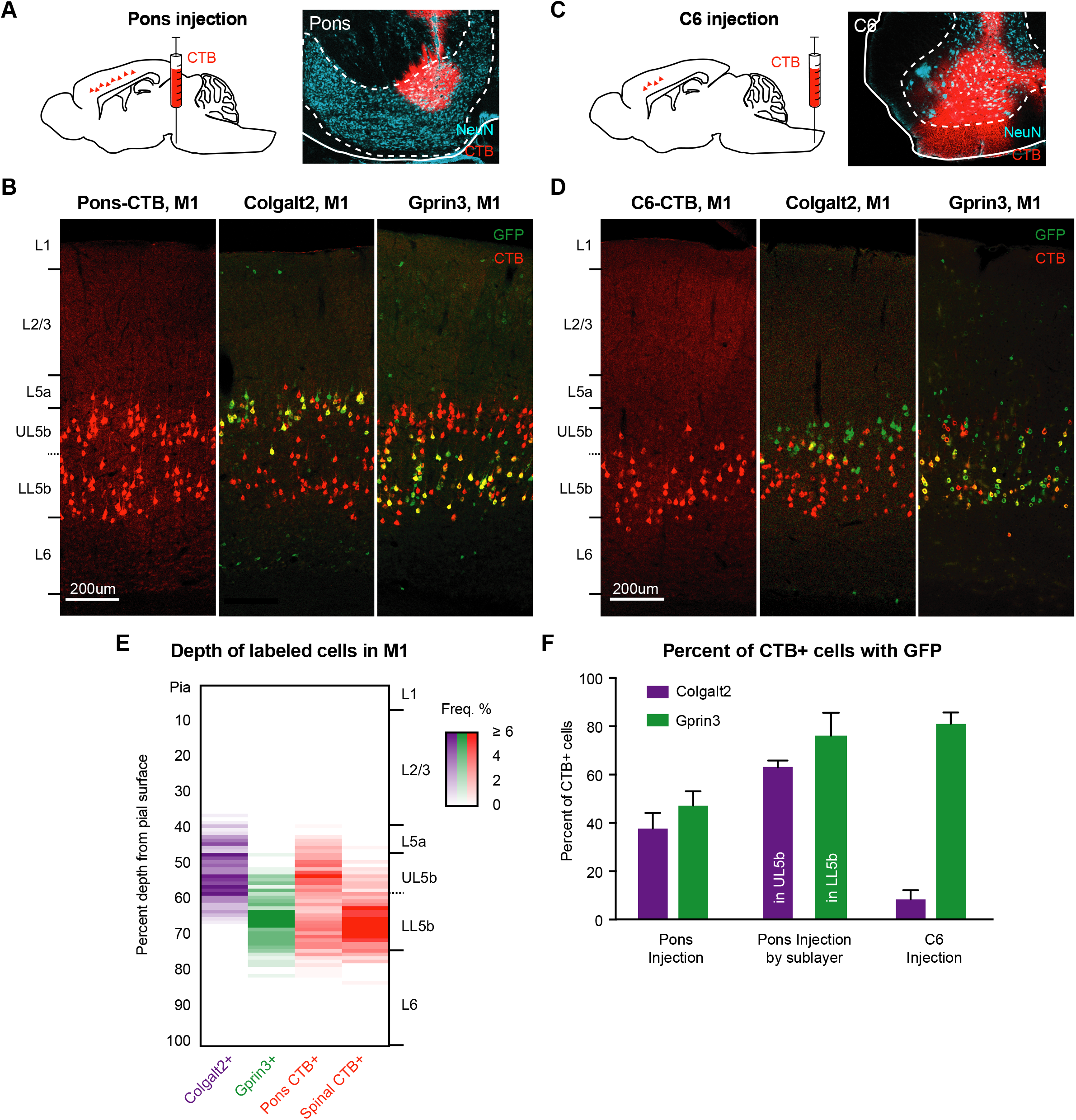
Retrograde tracing reveals overlapping and distinct axonal projection targets of Colgalt2 and Gprin3 cells. **A.** Schematic illustration of the strategy for retrograde tracing of corticopontine neurons with cholera toxin B (CTB) (left), and a representative image of the injection site (right) showing anti-CTB (red) and anti-NeuN (cyan) immunofluorescence. **B.** Immunofluorescent images showing anti-CTB labeled cells (red) in both sublayers of L5b in M1 following an injection into the pons (right). Staining with anti-GFP revealed overlap of labeled cells in UL5b of Colgalt2-bacTRAP animals (center) and LL5b in Gprin3-bacTRAP (right) animals. Scale bar = 200 μm. **C.** Schematic of the strategy for retrograde labeling of corticospinal neurons (left), and a representative image of the injection site in C6 spinal cord (right) showing anti-CTB (red) and anti-NeuN (cyan) immunoflueorscence. **D.** Immunofluorescent images, as in B, showing CTB-labeled cells in LL5b of M1 (left) with GFP-labeled cells in Gprin3-bacTRAP mice (right) but not Colgalt2-bacTRAP mice (center). **E.** Histogram of the quantification of laminar depth of GFP+ Colgalt2 (purple, n = 209 cells) and Gprin3 (green, n = 163) cells alongside CTB+ corticopontine (red left, n = 651 cells) and corticospinal cells (red right, n = 192 cells). Frequency reported as percent of total cells found at each depth. **F.** Quantification (mean + SEM) of the percent of M1 CTB+ cells that were GFP+ in Colgalt2-bacTRAP (purple) or Gprin3-bacTRAP (green) animals across all of L5b (left), or within each individual sublayer (center). Right shows the percent (mean + SEM) of double-labeled cells in M1 of each bacTRAP line following injections into C6.

In C6-injected animals, CTB+ cells were primarily observed in LL5b in M1, at a depth similar to Gprin3 neurons and distinctly below Colgalt2 cells (~60-75% from pia; **Figure 2D, E**). In Gprin3 bacTRAP brains, 82% of spinal CTB+ cells were EGFP+, while only 9% of spinal CTB+ cells were EGFP+ in Colgalt2 bacTRAP mice (**Figure 2F**). We also observed CTB co-localization with Gprin3 cells when CTB was injected into brainstem motor nuclei 7N and 5N (**Figure S3A-C**). Together, these data reveal that the subcortical targets of EGFP+ neurons in Colgalt2- and Gprin3-bacTRAP mice are consistent with PT projections and, while they both share a projection to the pons, the Gprin3 cells have additional distal collaterals that extend to the brainstem and cervical spinal cord.

### Gprin3 cells are vulnerable to degeneration in SOD1*G93A, while Colgalt2 neurons are resistant

Because Gprin3 cells showed a projection to the spinal cord that Colgalt2 cells lacked, we hypothesized that Gprin3 cells would be vulnerable to degeneration in a mouse model of ALS. To determine vulnerability of Colgalt2 and Gprin3 cells in disease, we crossed Colgalt2 and Gprin3 bacTRAP mice to hSOD1*G93A (B6JL.SOD1*G93A^+/−^) mice (**Figure 3A**), a widely used preclinical model of familial ALS (Chiu et al., 1995). SOD1*G93A-crossed mice began displaying tremors just after postnatal day 70 (P70), with severe deficits in rotarod performance appearing around eight weeks of age (**Figure S4A**). Colgalt2-bacTRAP::SOD1*G93A and Gprin3-bacTRAP::SOD1*G93A mice both reached end stage of disease at approximately P165, consistent with the typical survival time for this model (**Figure S4B**). We looked for signs of inflammation in M1 cortex of Colgalt2-bacTRAP::SOD1*G93A and Gprin3-bacTRAP::SOD1*G93A animals by staining for reactive GFAP+ astrocytes after P110 (**Figure 3B**). In WT animals, the majority of GFAP+ astrocytes were observed in deep L6 and in superficial L2/3 and L1 (**Figure 3B, C**). In SOD1*G93A animals, there was a marked increase in the number of GFAP+ astrocytes found in L5, between 40% and 75% depth from pia, suggesting the presence of dysfunction or degeneration in the vicinity of our Colgalt2 and Gprin3 neurons.

**Figure 3.**
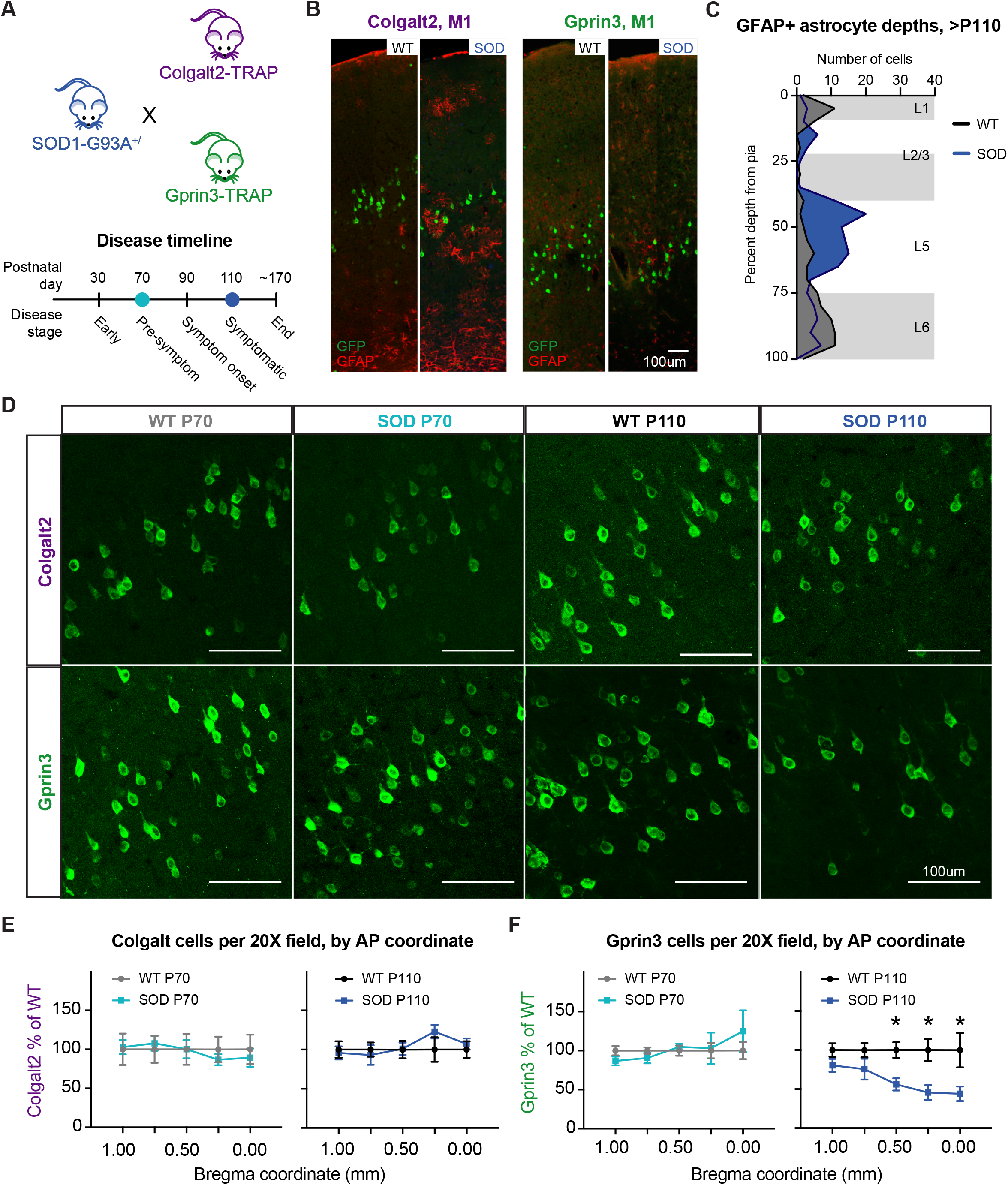
Loss of Gprin3 cells but not Colgalt2 cells in symptomatic SOD1*G93A mice. **A.** Schematic of breeding paradigm for SOD1*G93A experiments in Colgalt2-bacTRAP and Gprin3-bacTRAP animals (top), and the timeline of disease progression used for histology and sequencing (bottom). **B.** Immunofluorescent images of M1 from Colgalt2-bacTRAP and Gprin3-bacTRAP animals at the symptomatic time point, showing GFP (green) and the activated astrocyte marker, GFAP (red), across both healthy (WT) or disease (SOD) conditions. Scale bar = 100 μm. **C.** Frequency distribution of laminar depth of GFAP+ activated astrocytes across healthy (WT, grey, n = 83 cells) and disease (SOD, blue, n = 130 cells) conditions at age > P110. Mean depth of WT GFAP+ cells = 62%, SOD GFAP+ cells = 54% from pia. Frequency reported as absolute number of GFAP+ cells found at each depth. **D.** Representative images showing GFP+ cells in M1 of Colgalt2-bacTRAP (top row) or Gprin3-bacTRAP (bottom row) mice at pre-symptomatic (P70) and symptomatic (P110) time points in WT and symptomatic SOD animals. Scale bars = 100 μm. **E, F.** Quantification (mean ± SEM) of the number of GFP+ cells along the rostrocaudal axis of M1 at pre-symptomatic (left) and symptomatic (right) time points in healthy (WT, grays) and disease (SOD, blues) conditions for (**E**) the Colgalt2 bacTRAP line (WT P70 n = 5 animals, SOD P70 n = 8 animals, Colgalt2 WT P110 n = 7 animals, SOD P110 n = 5) and (**F**) the Gprin3 bacTRAP line (Gprin3 WT P70 n = 8 animals, SOD P70 n = 5 animals, Gprin3 WT P110 n = 7 animals, SOD P110 n = 5 animals). * p < 0.05 by unpaired t-test with Benjamini-Hochberg correction for multiple comparisons.

We next wanted to determine whether UL5b Colgalt2 or LL5b Gprin3 cells in M1 degenerate in SOD1*G93A animals. We counted the number of EGFP+ cells in serial coronal sections of M1 from Colgalt2-bacTRAP and Gprin3-bacTRAP mice expressing SOD1*G93A (SOD) or wildtype littermates (WT) to assess changes in cell number along the anterior-posterior extent of this region. Cells were counted across early symptomatic and late symptomatic time points, P70 and P110 respectively. We found no change in the number of Colgalt2 neurons along the extent of M1 (Bregma +1.00 mm to 0.00 mm) in SOD1*G93A mice (**Figure 3D, E**) at either age. While there was no difference in the number of Gprin3 neurons at P70, there was a significant decrease in the number of Gprin3 neurons in M1 at P110 (**Figures 3D** and **S4C**) with the greatest proportional loss found at more posterior coordinates (44% loss at Bregma +0.50 mm, and 55% at Bregma +0.25 mm; **Figure 3F**). This loss of Gprin3 neurons but not Colgalt2 neurons in SOD1*G93A mice reinforces the cell type-specificity of vulnerability even between closely related UL5b PT cells. These results also underscore the value of the Gprin3 and Colgalt2 bacTRAP lines which provide reproducible access to vulnerable and resilient populations, respectively.

### Vulnerable Gprin3 cells showed a robust molecular response to SOD1*G93A mutation

The vulnerability of LL5b Gprin3 cells and resistance of UL5b Colgalt2 cells in SOD1*G93A mice presented us with a unique opportunity to determine cell type-specific molecular responses of two closely related L5b populations during disease. To examine their gene expression patterns, we performed TRAP on SOD1*G93A mice and healthy littermates from both Colgalt2 and Gprin3 bacTRAP lines at a symptomatic age (~P110). Polysome-bound mRNAs were isolated following anti-EGFP immunoprecipitations (IP) on M1 homogenates and analyzed by RNA-seq along with transcripts extracted from whole tissue (input). We first confirmed the cell type specificity of TRAP isolations by determining relative expression of PT marker genes and glial genes in healthy and disease IP samples relative to cortical input samples (**Figure 4A**). All Colgalt2 and Gprin3 samples were enriched for classic PT cell type genes relative to cortical input, and were depleted for glial genes. Additionally, relative expression of these genes did not significantly change in disease, suggesting that these cell type markers are robust even in an ALS context.

**Figure 4.**
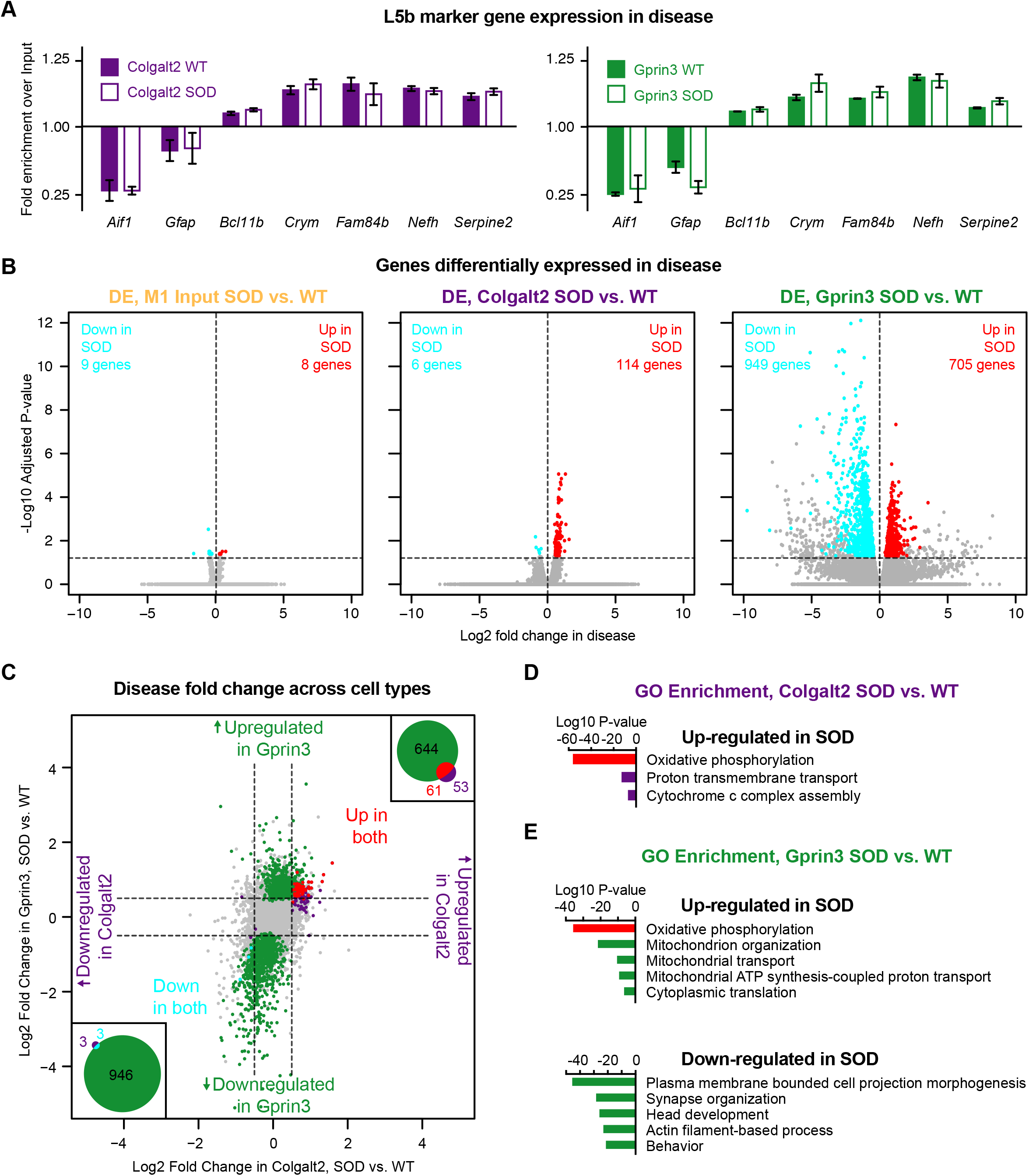
Gprin3 cells show a robust molecular response to SOD1*G93A expression. **A.** Bar graphs showing relative expression (mean ± SD CPM) of glial and L5b marker genes across healthy (WT) and disease (SOD) TRAP samples from Colgalt2 and Gprin3 cells compared to sample-matched M1 input. **B.** Volcano plots identifying DE genes between WT and SOD littermates for M1 input (left), Colgalt2 TRAP (center), and Gprin3 TRAP (right) samples. Genes significantly upregulated (red) or downregulated (cyan) in SOD are indicated. Dotted lines represent significance thresholds of 0.0 log_2_ fold change (vertical line) and 1.3 −log_10_ padj (horizontal line). **C.** Scatterplot comparing SOD-mediated log_2_ fold changes in Colgalt2 TRAP (x-axis) and Gprin3 TRAP (y-axis) samples for all genes that had mean CPM > 100 in Gprin3 cells. Purple and green dots highlight genes that were significantly changed only in Colgalt2 cells or Gprin3 cells respectively while genes that were changed in both cell types are highlighted in red (upregulated) or cyan (downregulated). Dotted lines represent log_2_ fold change of +/− 0.5. Insets show Venn diagrams indicating the total number of genes that were uniquely or commonly upregulated (top right) or downregulated (bottom left) across both cell types. **D.** Summary of GO enrichment analysis of DE genes in Colgalt2 TRAP data from B. **E.** Summary of GO enrichment analysis of DE genes in Gprin3 TRAP data from B. Red bars in D and E indicate a category that was common to both cell types.

We next wanted to identify genes that showed a significant change in expression in disease by performing differential gene expression (DE) analysis between healthy (WT) and disease (SOD) samples. DE revealed that only 17 genes were significantly regulated (padj < 0.05 and baseMean > 100 log_10_CPM) in M1 whole tissue (input) samples in response to SOD1*G93A expression (**Figures 4B** and **S5A**). Resistant Colgalt2 cells instead showed 120 genes with significant changes in expression in disease, with 114 up-regulated and six genes down-regulated in the SOD condition (**Figures 4B** and **S5A**). Vulnerable Gprin3 cells, in contrast, showed the most robust response to SOD1*G93A expression, with 1,654 genes differentially regulated (705 up-regulated and 949 down-regulated). To identify genes commonly dysregulated in both Gprin3 and Colgalt2 cells, we compared the fold-change in disease for each gene across the two cell types and found 61 genes were significantly up-regulated in both cell types while only three were commonly down-regulated (**Figures 4C** and **S5B**). Lastly, to determine the biological and cellular roles of disease-modulated genes, we performed gene ontology (GO) analyses on all DE genes from Colgalt2 or Gprin3 cells (**Figures 4D, E** and **S5C**). Genes related to Oxphos were up-regulated in both cell types, while Gprin3 cells additionally up-regulated genes associated with translation and mitophagy (**Figure 4E** and **S5C**). Genes associated with regulation and maintenance of neuronal processes- i.e. axons, dendrites, and synapses- were down-regulated in Gprin3 cells. These analyses revealed that vulnerable Gprin3 cells had a relatively robust molecular response to SOD1*G93A. They also highlight molecular changes that occur as a general response to the presence of SOD1*G93A in PT cells, while at the same time reveal potential cell type-specific adaptations in the vulnerable LL5b PT cell type.

### Comparing Colgalt2 and Gprin3 gene expression changes reveal cell type-specific molecular responses to SOD1*G93A

Vulnerable Gprin3 and resistant Colgalt2 cells showed a subset of common changes in gene expression, but Gprin3 cells showed additional and robust changes associated with other biological pathways. To understand the potential cell type-specificity of these molecular responses, we determined if individual genes that compose these pathways were indeed uniquely changed only in Gprin3 cells, or if Colgalt2 cells showed potentially similar but non-significant changes in these genes with expression of SOD1*G93A. Synaptic and axonal genes that were down-regulated in Gprin3 cells showed almost no change in expression between SOD and WT in M1 inputs or Colgalt2 TRAP data while they decreased by 50% in Gprin3 TRAP data (**Figure 5A, B**). Therefore, these genes are uniquely modulated in the ALS-vulnerable Gprin3 cell type. Since cytoplasmic translation GO term was upregulated with disease in Gprin3 cells (**Figure 4E**), we next looked at ribosomal genes across our two cell types. Surprisingly, both Gprin3 and Colgalt2 cells showed positive fold-changes for almost all small and large ribosomal subunit genes, though Gprin3 cells showed a greater increase relative to Colgalt2 cells (**Figure 5C, D**). Because mitochondrial genes appeared to be strongly up-regulated in both Colgalt2 and Gprin3 cells in ALS, we also looked at changes in expression of mitochondrial ribosomes across our two cell types (**Figure 5C, D**). Once again, we observed that both Colgalt2 and Gprin3 cells appeared to increase expression of these translation-associated genes. In fact, Colgalt2 cells showed a 1.3-fold mean increase in both cytoplasmic and mitochondrial ribosomal genes expression, and Gprin3 cells showed a significant 1.49-fold mean increase for cytoplasmic ribosomes and a 1.41-fold mean increase for mitochondrial ribosomal genes (**Figure 5D**). There was no change in these genes in M1 input suggesting that this may be an adaptation specific to PT cells in response to SOD1*G93A. Additionally, these data reveal that the pathways selectively regulated in Gprin3 cells, such as synaptic and axonal structure and function, may underlie upper motor neuron-specific pathology in the disease, and contribute to selective loss of these vulnerable cells.

**Figure 5.**
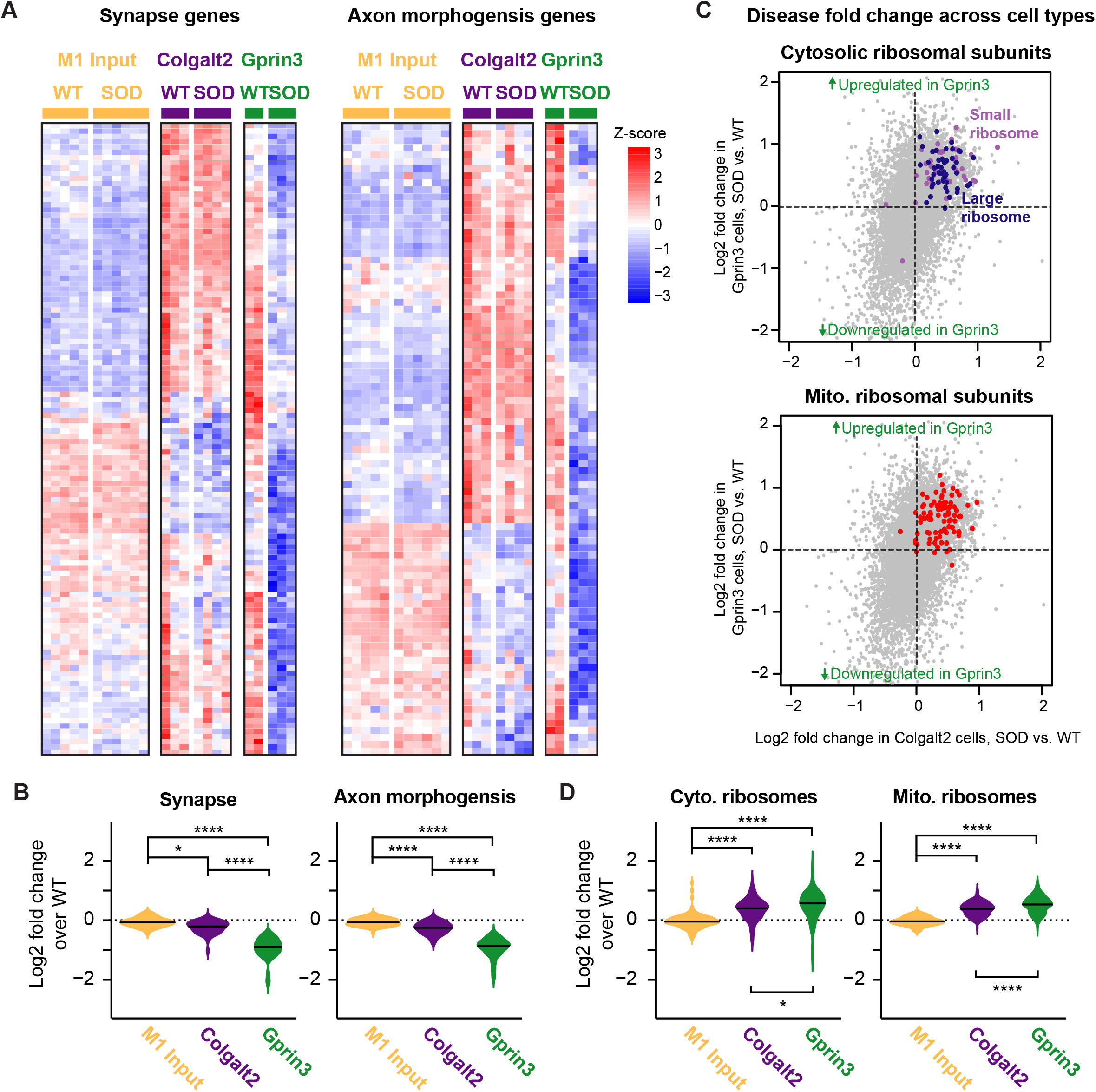
Gprin3 cells modulate ribosomal, axon morphogenesis, and synapse structure and function genes in disease. **A.** Heatmaps showing relative expression of genes that code for synaptic proteins (left) and axon morphogenesis proteins (right) in M1 input, Colgalt2 TRAP, and Gprin3 TRAP samples across WT and SOD replicates. Values reported as z-scores for normalized CPM, scaled for each gene. **B.** Violin plots of SOD-mediated log_2_ fold change values for all synapse (left) and axonal (right) genes that showed a significant change in Gprin3 cells across M1 input, Colgalt2, and Gprin3 cells. **C.** Scatterplot showing SOD-mediated log_2_ fold changes in Colgalt2 (x-axis) and Gprin3 (y-axis) TRAP samples for cytosolic ribosomal subunit genes (top, blue and lilac dots) and mitochondrial ribosome genes (bottom, red dots) among all genes that had mean CPM > 100 in Gprin3 cells. **D.** Violin plots of SOD-mediated log_2_ fold change values of cytoplasmic (left) and mitochondrial (right) ribosome genes from C across M1 input, Colgalt2 TRAP, and Gprin3 TRAP samples. For panels B and D, * p-value < 0.05 and **** p-value < 0.00001 by one-way ANOVA and subsequent Tukey multiple comparison test.

### Oxphos genes were up-regulated across both resistant Colgalt2 and vulnerable Gprin3 cells, but show differential baseline levels of expression between these two cell types

One set of genes that showed a significant up-regulation by DE in both Colgalt2 and Gprin3 cells, belonged to the Oxphos pathway (**Figure 4C, D**). This increase suggested that a change in respiration may be a default response to SOD1*G93A expression in L5b cells. We first wanted to determine whether this up-regulation of Oxphos genes included all complexes of the electron transport chain (ETC) or was selective to specific subunits. We compared the relative expression level of each gene composing the individual complexes of Oxphos (**Figure 6A, B**), and found that Colgalt2 and Gprin3 cells showed a significant up-regulation of all functional complexes by an average of 1.42- and 1.41-fold respectively in SOD, while M1 input samples did not show any change in these groups (**Figures 6B** and **S6B, C**). We also noted a small but non-significant up-regulation of genes that compose the tri-carboxylic acid cycle (TCA) in both Colgalt2 and Gprin3 cells in disease (**Figure S6B**), and no changes in genes coding for the glycolysis pathway (**Figure 6B**), suggesting that while aerobic metabolism may be increased in disease, anaerobic metabolism does not appear to be modulated. Gprin3 cells exhibited a unique up-regulation of genes involved in mitochondrial transmembrane transport (1.27-fold average increase) in SOD animals, and a greater increase in a subset of mitochondrial autophagy genes (1.15-fold in Colgalt2 cells, 1.61-fold in Gprin3 cells on average), suggesting that while an increase in respiration may be a common response to SOD1*G93A expression across L5b cells, certain important mitochondria-associated mechanisms are more strongly modulated only in vulnerable LL5b cells.

**Figure 6.**
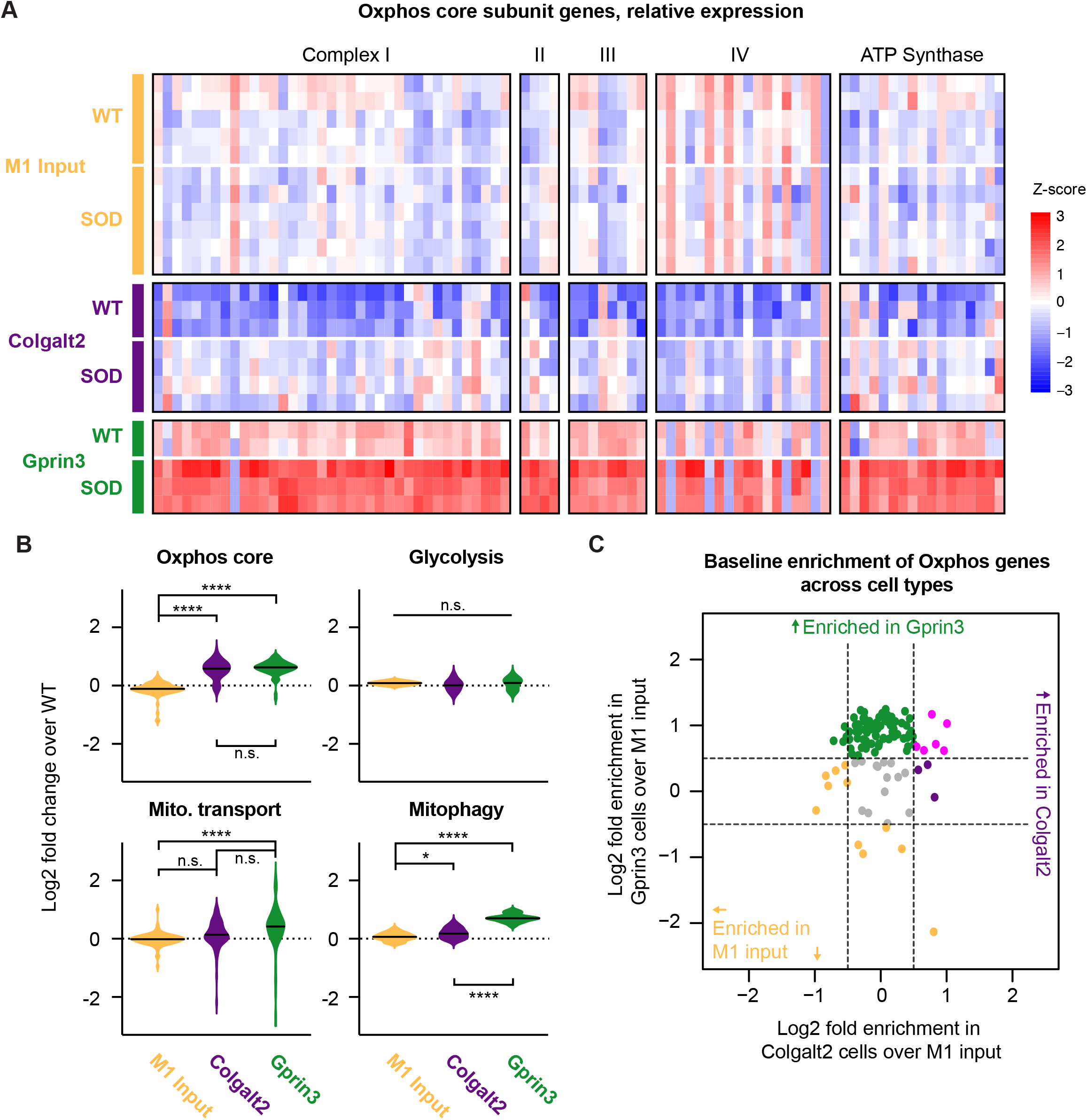
SOD1*G93A-mediated regulation of oxidative phosphorylation in layer 5b PT neurons. **A.**Heatmap showing relative expression of genes encoding each subunit of the Oxphos complexes in each experimental condition. Values reported as z-scores for normalized CPM, scaled for each gene. Violin plots of SOD-mediated log_2_ fold change values for all Oxphos core subunit and glycolysis genes (top), as well as all mitochondrial transmembrane transport and mitophagy genes (bottom) across M1 input, Colgalt2 TRAP, and Gprin3 TRAP samples. * p-value < 0.05, **** p-value < 1 × 10^−7^, n.s. not significant by one-way ANOVA and subsequent Tukey multiple comparison test. **C.** Scatterplot comparing the enrichment of Oxphos genes in Colgalt2 (x-axis) and Gprin3 (y-axis) at baseline. Genes enriched in Gprin3 only (green), Colgalt2 only (purple), both L5b cell types (magenta), or depleted from each cell type relative to M1 input (orange) are indicated.

While both Colgalt2 and Gprin3 cells exhibited a significant upregulation in Oxphos-related genes in SOD1*G93A mice, these genes are expressed at higher levels in WT Gprin3 cells compared to WT Colgalt2 cells (**Figures 6A** and **S6B, C**). We therefore assessed expression levels of Oxphos genes across Colgalt2 and Gprin3 cells relative to M1 input using baseline DE TRAP data, e.g. not crossed to the SOD1*93A mice (**Figures 1E** and **S2B**). Gprin3 cells had a median 1.75-fold higher level of Oxphos gene expression than Colgalt2 cells (**Figure 6C**) and oxidative phosphorylation is the most enriched GO category in Gprin3 DE genes compared to both M1 input and Colgalt2 cells (**Figure S2C, E**). This suggests that Gprin3 cells not only increase their levels of Oxphos gene expression in response to SOD1*G93A, but that they inherently express higher these genes at higher levels compared to Colgalt2 neurons.

### Expression pattern of a hypoxia- and antioxidant-responsive transcriptional program underlies cell type-specific changes in vulnerable Gprin3 cells

We hypothesized that a disease-related increase in Oxphos output over the already elevated level of respiration in Gprin3 cells at baseline could correspondingly increase production of reactive oxygen species (ROS) and increase cell stress (**Figure 7A**). To test this, we examined whether the expression levels of genes in ROS response pathways were significantly modulated in these cells in response to SOD1*G93A. In Gprin3 cells, a set of 45 oxidative stress and antioxidant genes showed significant changes in disease, with 20 showing an up-regulation and 25 showing a down-regulation (**Figure 7B**). Similar changes were not observed in the whole-tissue M1 input samples, and only four of these genes- *Ndufa12*, *Romo1*, *Selenok*, and *Sod1*- were up-regulated in Colgalt2 cells as well. Notably, nine of the down-regulated genes in Gprin3 cells– including *Hif1a*, *Epas1*, and *Nfe2l2*– code for transcription factors important for controlling cellular responses to hypoxia and oxidative stress conditions (Hamanaka and Chandel, 2009; Kobayashi et al., 2009; Semenza, 2001). Examination of the relative expression levels of these and other hypoxia and oxidative stress response factors across baseline Gprin3 and Colgalt2 data revealed that Gprin3 cells innately expressed lower levels of the factors *Foxo1*, *Epas1*, *Nfe2l2*, and *Notch1* compared to Colgalt2 cells and M1 input (**Figures 7C** and **S7B**), with *Epas1* and *Notch1* showing an ~11.5- and 4.5-fold enrichment in Colgalt2 cells over Gprin3 cells respectively. Together these data highlight the fact that while Gprin3 cells appear to decrease expression of these oxygen species-reactive factors under conditions of increased Oxphos, these cells show lower levels of these factors even at healthy baseline. This suggests that in Gprin3 cells, the underlying expression pattern of the oxygen species-responsive transcriptional program, compounded with a down-regulation during disease, may have severe consequences for the ability of Gprin3 cells to manage increased ROS production in the presences of SOD1*G93A.

**Figure 7.**
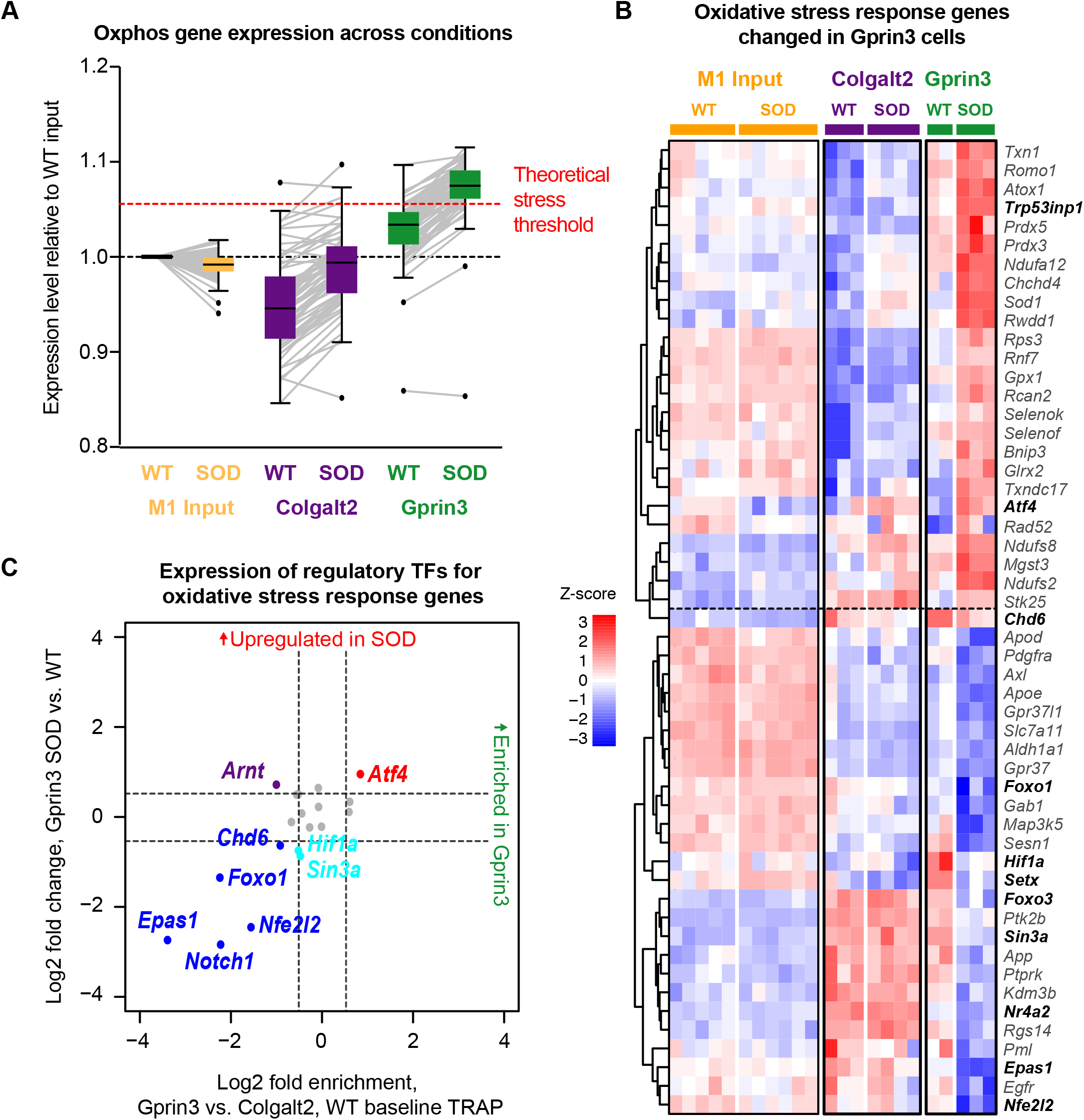
Oxidative stress response genes are modulated in Gprin3 cells, and hypoxia- and oxidative stress-responsive transcription factors are depleted in healthy Gprin3 cells. **A.** Box and whisker plot showing ratio of CPMs of genes that comprise the Oxphos core subunits relative to WT M1 input for WT and SOD data sets across M1 input, Colgalt2 TRAP, and Gprin3 TRAP samples. Gray lines indicate trajectory of change for each individual Oxphos gene. Dotted red line highlights a theoretical threshold above which Oxphos gene expression levels may trigger oxidative stress pathways. **B.** Heatmap showing relative expression across all samples of genes involved in oxidative stress and antioxidant responses that were significantly changed in Gprin3 cell. Genes that code for transcription factors are highlighted in bolded text. Values reported as z-scores for normalized CPM, scaled for each gene. **C.** Scatterplot comparing enrichment of genes encoding transcription factors associated with activation of hypoxia and oxidative stress response pathways between Gprin3 and. Colgalt2 at baseline (x-axis) and Gprin3 cells in disease (y-axis). Genes significantly depleted in Gprin3 compared to Colgalt2 and significantly down-regulated in disease are highlighted in blue, genes down-regulated in disease but not enriched in either cell type are highlighted in cyan, genes depleted in Gprin3 cells but not changed in disease are highlighted in purple, and genes that were up-regulated in disease but not enriched in either cell type are highlighted in red.

## DISCUSSION

The selective vulnerability of corticospinal L5b PT neurons in the primary motor cortex (M1) to ALS-causing mutations has been described across species (Hammer et al., 1979; Ozdinler et al., 2011; Thomsen et al., 2014; Zang and Cheema, 2002), although the mechanisms that underlie their degeneration compared to other L5b cell types have remained elusive. Using TRAP molecular profiling and detailed anatomy, we show that Gprin3-bacTRAP mice labeled a population of corticospinal projection neurons located in lower L5b of motor cortex that degenerated in SOD1*G93A mice. These cells showed a robust molecular response in disease compared to resilient upper L5b corticopontine cells labeled by Colgalt2-bacTRAP mice and the rest of motor cortex. Analysis of the TRAP data revealed that while both PT populations up-regulated pathways involved in mitochondrial function and oxidative phosphorylation, elevated expression of metabolic genes at baseline and a deficiency in antioxidant signaling in disease underlie selective degeneration of the Gprin3 cells. These results demonstrate that while there is overlap in the cellular adaptations by vulnerable and resilient cell types to disease-causing mutations, their intrinsic properties dictate how the cell responds to these adaptations to determine survival or degeneration and may provide a molecular definition of vulnerability to disease.

M1 is important for motor planning, learning, and execution and is widely connected throughout the CNS through the long distance projections of L5b PT cells (Gu et al., 2017; Guo et al., 2015; Kawai et al., 2015; Suter et al., 2013). The repertoire of genes expressed by Gprin3 and Colgalt2 cells were highly correlated with each other relative to other neuronal types in the cortex, driven by genes associated with L5b pyramidal neuron identity. Within M1, the distribution of Colgalt2 and Gprin3 cells was divided into UL5b and LL5b, respectively, but both cell populations converged into the same layer in other regions of cortex. This is consistent with the variation of the thickness of cortical laminae across the different functional areas of cortex, with an expansion of L5b in the motor regions of cortex (Caviness, 1975; Cederquist et al., 2013). Further, local intracortical afferents have been shown to vary between upper and lower subdivisions of L5b in M1 (Hooks et al., 2013) while single cell analysis identified genetically distinct pyramidal cell populations populating these sublayers in anterolateral motor cortex (ALM; (Economo et al., 2018; Tasic et al., 2018). The divergence of axonal projections of Colgalt2 and Gprin3 cells imply unique roles for each cell type in motor output and corresponds to the distinct projection patterns of UL5b and LL5b in ALM (Economo et al., 2018). Previously generated transgenic mouse lines that target L5b, such as the commonly used Thy1-EYFP line (Bareyre et al., 2005; Feng et al., 2000; Yu et al., 2008) and Uchl1-EGFP line (Yasvoina et al., 2013), show expression of reporters and constructs across all of L5b rather than in its distinct sublayers. Therefore, our use of BAC transgenics enabled reproducible molecular and anatomic access to both vulnerable and resilient cell populations across disease phenotypes, presenting us with a rare opportunity to compare disease-relevant molecular differences between two closely related neuron types.

TRAP profiling revealed significant molecular adaptations in both PT populations in symptomatic SOD1*G93A mice. While this response was much more robust in the vulnerable Gprin3 cells, a number of these changes were observed to a lesser degree in Colgalt2 neurons, highlighting a commonality of cellular responses to SOD1*G93A expression in both L5b PT cell types. Most notably, both cells exhibited an up-regulation of mitochondrial genes, specifically those involved in Oxphos pathways. Studies from human patients and animal models have shown that mitochondrial dysfunction is a common, early phenomenon in ALS and other neurodegenerative diseases. Neuronal survival is critically dependent on the integrity and functionality of mitochondria. Mitochondria are the primary site for the generation of ATP through Oxphos– accounting for 90% of all ATP generated in the CNS (Hyder et al., 2013)– and as a byproduct are the major source of ROS (Adam-Vizi and Chinopoulos, 2006; Hirst et al., 2008; Kudin et al., 2004; Lenaz et al., 2002; Muller et al., 2004; Nunnari and Suomalainen, 2012). The appearance of deficits in Oxphos, mitochondrial calcium buffering, generation of ROS, and mitochondrial transport have been observed prior to the onset of ALS symptoms (Bowling et al., 1993; Browne et al., 2006; Jaiswal, 2014; Sasaki et al., 2005), and aggregation of mutant SOD1 within mitochondria leads to free radical generation, ETC disruption, loss of mitochondrial membrane potential, diminished Oxphos and decreased ATP production (Mattiazzi et al., 2002; Vandoorne et al., 2018). While most evidence for altered mitochondrial function has come from spinal cord or cultured motor neurons, high throughput sequencing of frontal cortex from patients with sporadic or C9ORF72-linked ALS found Oxphos genes were dyregulated in both cases (Prudencio et al., 2015). Oxidative stress, including an up-regulation of Oxphos, was also a defining feature of a subset of motor cortex samples from a separate cohort of ALS patients (Tam et al., 2019). Since we did not detect an up-regulation of Oxphos in our M1 whole tissue input samples, it is likely that the modulation of aerobic metabolism is a specific adaptation of L5b PT cells rather than a global response to the disease-causing mutation.

The unique responses of the vulnerable Gprin3 cells reflected adaptations that indicate cellular stress and neurodegenerative mechanisms, including an up-regulation of genes associated with mitophagy, which was not seen in the resilient Colgalt2 population. Mitophagy refers to the selective removal of dysfunctional mitochondria and is a key strategy for mitigating cellular stress, and can even be used to delay the initiation of cell death (Green et al., 2011; Twig and Shirihai, 2011). Prompt removal of damaged mitochondria is critical for cell viability and the energy demands of neurons. Inefficient turnover of mitochondria by ALS-associated mutations in *TBK1* and *OPTN genes*, highlights a potential role for impaired mitophagy in ALS (Evans and Holzbaur, 2019; Moore and Holzbaur, 2016; Wong and Holzbaur, 2014). An increase in mitophagic gene expression likely reflects a pro-survival adaptation in Gprin3 cells occurring at the symptomatic stage. Gprin3 cells also down-regulated genes involved in maintaining synaptic and axonal structure and function, suggesting that these cells may undergo morphological changes associated with dysfunction and cell death. Dendritic and synaptic atrophy in L5b neurons in M1 have been previously reported, even at a pre-symptomatic age (Fogarty et al., 2015; Saba et al., 2015), which may include a down-regulation of synaptic transmission genes (Kim et al., 2017). Our findings suggest that these changes may persist beyond this pre-symptomatic stage and are most robust in the vulnerable corticospinal subpopulation of L5b neurons. Additionally, cytoskeleton-dependent axon transport defects have been previously reported in spinal motor neurons in SOD1*G93A animals (Warita et al., 1999; Williamson and Cleveland, 1999; Zhang et al., 1997). The decrease in axon morphogenesis-associated gene expression that we observed suggests this phenomenon may also occur in vulnerable upper motor neurons.

The neurodegeneration of Gprin3 cells and resilience of Colgalt2 cells to SOD1*G93A expression, despite their substantial overlap in cellular adaptations in symptomatic animals, raises the possibility that the inherent molecular properties of these cells drive divergent consequences in disease. In healthy animals, Gprin3 cells had higher baseline levels of Oxphos gene expression than Colgalt2 cells and total M1 tissue. This suggests that corticospinal neurons may have increased bioenergetic demands compared to other neurons in M1. Little is known about how metabolic demands and/or output varies between neuronal types. It is possible that Gprin3 cells require more ATP production in order to maintain their increased axonal volume due to long-range projections and extensive collaterals typical of corticospinal neurons. Indeed, metabolic demand has been linked to cell size in photoreceptors and surface area of the cell membrane (Le Masson et al., 2014; Niven et al., 2007; Sengupta et al., 2010). In lower motor neurons of the spinal cord, greater vulnerability appears to positively correlate with cell size (Pun et al., 2006; Robberecht and Philips, 2013). Fast fatigable alpha motor neurons have larger soma size, axon diameter, more dendritic branching, and greater innervation fields when compared to ALS resilient fast fatigue-resistant alpha motor neurons or slow gamma motor neurons (Ragagnin et al., 2019; Shoenfeld et al., 2014). This supports the notion that larger neurons may have greater metabolic needs, consequently making them particularly vulnerable to energetic challenges arising in ALS.

Surprisingly, even though Gprin3 neurons show higher levels of Oxphos gene expression, respiration rates have been shown to decrease in ALS even before pathologies develop (Browne et al., 2006; Cozzolino and Carri, 2012; Irvin et al., 2015; Kawamata and Manfredi, 2010; Peixoto et al., 2013). Although this evidence comes largely from in vitro studies, this trend is apparent across both SOD1*G93A mice (Irvin et al., 2015) and clinical patients (Ghiasi et al., 2012). Despite increased Oxphos gene expression acting as a compensatory mechanism to restabilize homeostasis, Gprin3 cells likely fail to meet energetic demands caused by a stressor such as SOD1*G93A. Follow up studies are necessary to tease out the source of the metabolic instability unique to Gprin3 mitochondria. Interestingly, even though they show higher levels of Oxphos gene expression, we found that transcription factors downstream of mitochondrial ROS are expressed at lower levels in Gprin3 cells. These factors are critical regulators of the cell’s response to toxic ROS (Hamanaka and Chandel, 2009; Kobayashi et al., 2009), and deletion of these factors leads to cellular pathologies (Lee et al., 2003a; Lee et al., 2003b; Scortegagna et al., 2003). One possible explanation for the selective vulnerability of Gprin3 cells may be that despite increased Oxphos, they are poorly equipped to handle changes in the balance of oxygen species. This phenomenon has been demonstrated in cerebellar granule and hippocampal CA1 neurons which are inherently more sensitive to oxidative damage (Wang and Michaelis, 2010; Wang et al., 2009). In particular, despite their close proximity and similar cell morphology, CA1 neurons show selective vulnerability to oxidative stress, while CA3 are spared (Wang et al., 2005). The unusually high energy demands of motor neurons may be largely met by mitochondria, with the consequence of increased production of ROS. High ROS production by motor neurons may shed light on why SOD1, which is ubiquitously expressed, is particularly abundant in these cells (Pardo et al., 1995). Hence, it is possible that similar to other oxidative stress-sensitive neurons, an inherently high baseline of ROS may make Gprin3 cells more vulnerable to further increases in ROS production. In a situation where the cells enter a state of acute oxidative stress or hypoxia, Gprin3 cells may be slower to respond than neighboring cells. It is therefore paradoxical that with an up-regulation of Oxphos genes in disease, Gprin3 cells further down-regulate the expression of these antioxidant factors. Future work should more directly test how these factors affect the sensitivity of L5b neurons to ALS-causing mutations in order to understand how a neuron’s ability to regulate levels of respiration and ROS may determine its vulnerability in ALS.

By comparing two highly related yet distinct L5b projection neuron populations that have differential vulnerability to degeneration in a preclinical ALS model, we show that a defining feature of a vulnerable population is an elevated expression of genes involved in energy production and that a further up-regulation of this pathway is a key molecular signature for cellular responses to disease. This is one of the first examples of a specific, intrinsic cellular property of a vulnerable population that is also uniquely, and directly, altered by disease pathology in ALS. Further studies are needed to flesh out the link between cell type specific differences in the expression of nuclear encoded mitochondrial genes, energetic output, and neurodegeneration.

## Supporting information

Moya_Supplementary_Material

## ACKNOWLEDGEMENTS

This work was supported by NIH/NINDS grants R01NS091722 (E.F.S.) and R21NS105047 (E.F.S.), ALS Therapy Alliance Grant 2013-F-052 (E.F.S.), ALS Association grants 14DUYT and 1114-471-454 (E.F.S.), and the National Science Foundation Graduate Research Fellowship 2014165948 (M.V.M.). N.H. is an investigator of the Howard Hughes Medical Institute (HHMI). We thank The Rockefeller University Genomics Resource Center, Transgenic Services Laboratory, and Comparative Bioscience Center as well as the New York Genome Center for technical support. We also thank Dr. Thomas Carroll for bioinformatics support as well as Drs. Hemali Phatnani and Ines Ibanez-Tallon for critical discussions.

## AUTHOR CONTRIBUTIONS

M.V.M, N.H., and E.F.S. conceived and designed the study. M.V.M., E.F.S., R.K., M.N.R., B.A.C., S.B.P. and C.E.S. performed all experiments. M.V.M. and E.F.S. analyzed data. M.V.M. and E.F.S. wrote the manuscript with comments from B.A.C., R.K., and N.H.

