## Supplementary material for "Unique molecular features and cellular responses differentiate two populations of motor cortical layer 5b neurons in a preclinical model of ALS": Moya_Supplementary_Material

### *Supplemental Information*

Figures S1-S7 begin on next page.

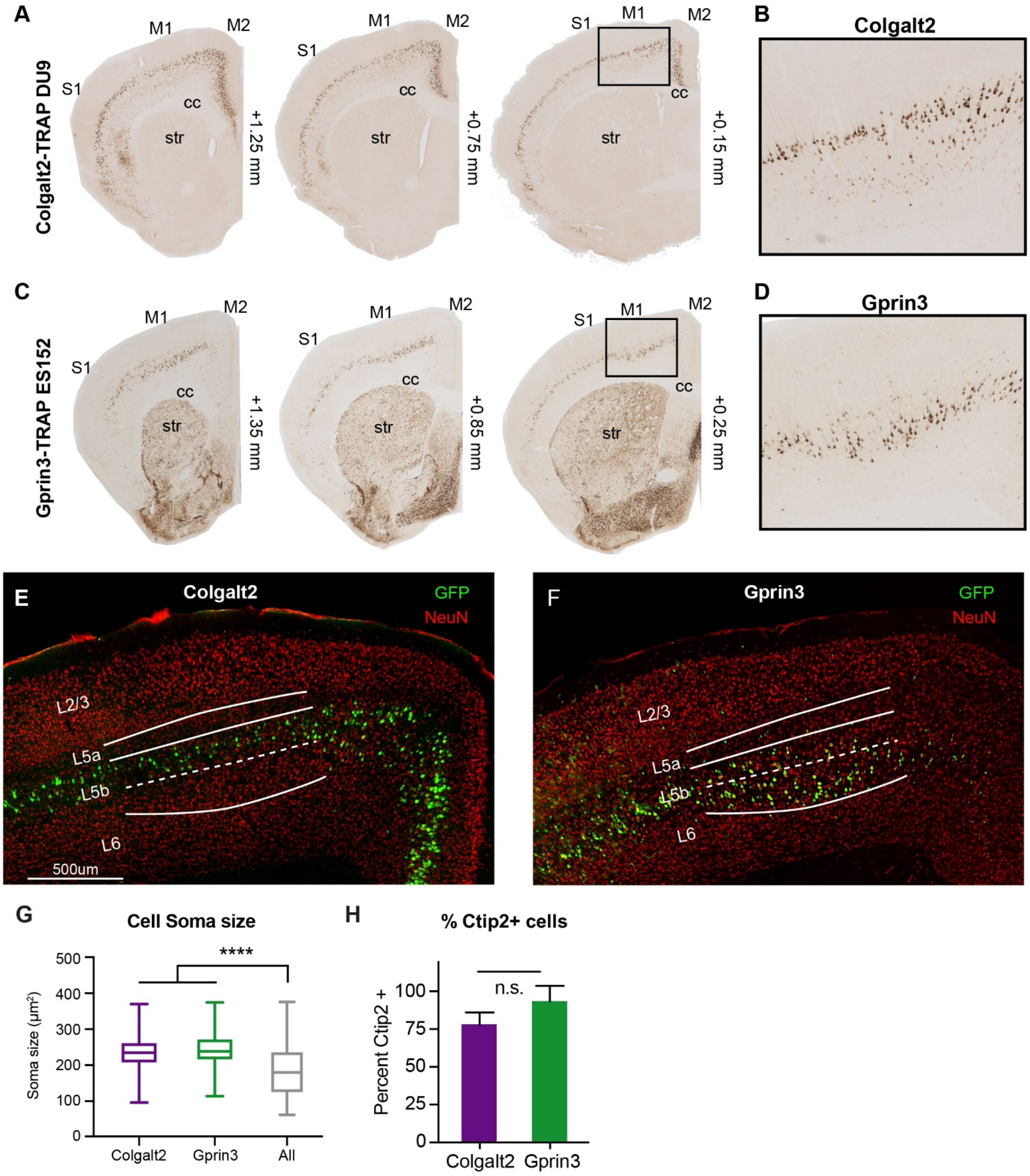


**Figure S1. Colgalt2-TRAP DU9 and Gprin3-TRAP ES152 mice respectively target two laminarly distinct L5b pyramidal neurons, related to Figure 1.** A) DAB immunostaining shows EGFP+ cells across the anterior-posterior (AP) extent of M1 in a Colgalt2-TRAP mouse. S1 = primary somatosensory cortex, M1 = primary motor cortex, M2 = secondary motor cortex, str = striatum, cc =

corpus callosum. B) Inset shows upper L5b (UL5b) localization of EGFP+ Colgalt2 cells. C) Similar staining and AP coordinates as in A, showing the distribution of EGFP+ cells in a Gprin3-TRAP animal. D) Inset showing lower L5b (LL5b) localization of EGFP+ Gprin3 cells. E and F) Fluorescent immunostaining showing distinct L5b laminar distribution of EGFP+ cells in Colgalt2-TRAP (E) and Gprin3-TRAP (F) animals. Red channel shows NeuN+ cells, green channel shows EGFP+ TRAP cells. Scale bar = 500 µm. G) Box and whisker plots showing population distribution for soma sizes of Colgalt2 cells (purple, n = 292 cells), Gprin3 cells (green, n = 285 cells), and randomly selected M1 pyramidal neurons (“All”, grey, n = 1253 cells). Values reported as $\mu m^2$. **** p-value < 0.0001 by one-way ANOVA and subsequent Tukey multiple comparison test. H) Co-localization percentage of Ctip2 with EGFP+ Colgalt2 cells (purple bar, 151 Colgalt2+/Ctip2+ cells out of n = 191 Colgalt2 cells), or Gprin3 cells (green bar, 127 Gprin3+/Ctip2+ out of n = 135 Gprin3 cells). n.s. = not significant by two-tailed t-test.

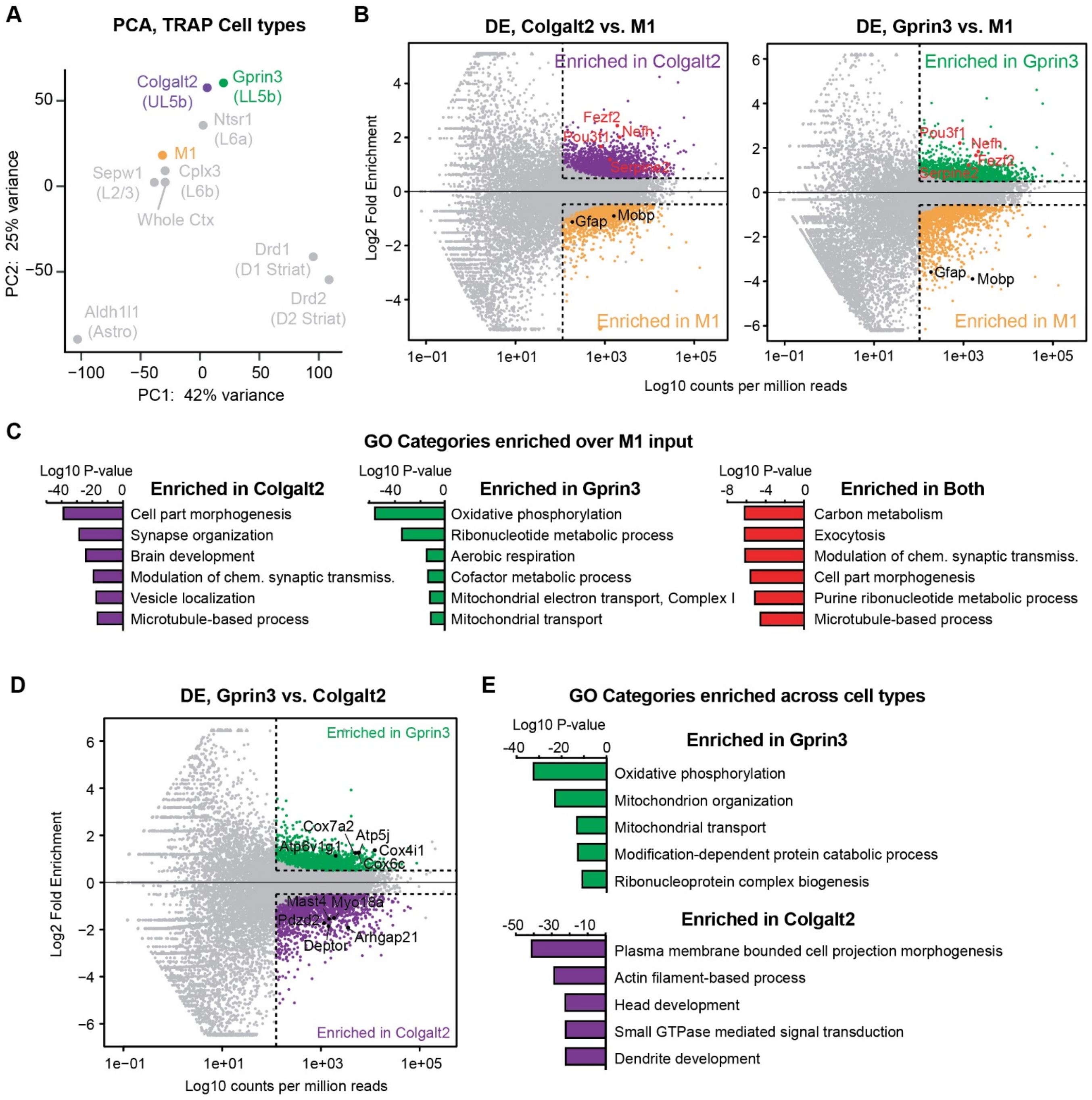


**Figure S2. TRAP sequencing reveals differential gene expression patterns from Colgalt2 vs. M1 input, Gprin3 vs. M1 input, and Gprin3 vs. Colgalt2 comparisons, related to Figure 1.** A) PCA plot shows whole transcriptome mapping along the first 2 components for various cell types in the cortex, including M1 input (orange), Colgalt2 (purple), and Gprin3 (green) cells. B) MA plots of differential expression between Colgalt2 and M1 input (left panel), and Gprin3 and M1 input (right panel). Genes that were significantly (adjusted p-value $< 0.05$ and mean CPM $> 100$) enriched in Colgalt2 cells are shown in purple, genes that were enriched in Gprin3 cells are shown in green, and genes that were enriched in M1 input are shown in orange. Known L5b cell type markers and glial genes are labeled. C)

Genes that were significantly enriched in Colgalt2 cells (purple), in Gprin3 cells (green), or in both cell types (red) relative to M1 input were run through Metascape functional GO analysis, and adjusted p-values for each resulting category are shown. D) MA plot showing differential expression between Gprin3 and Colgalt2 cells, with genes significantly (adjusted p-value < 0.05 and mean CPM > 100) enriched in Gprin3 cells (green) and significantly enriched in Colgalt2 cells (purple) highlighted. A subset of these genes is labeled. E) Enriched GO categories for differentially enriched genes in Gprin3 (green bars) and Colgalt2 cells (purple bars). Values shown are adjusted p-values.

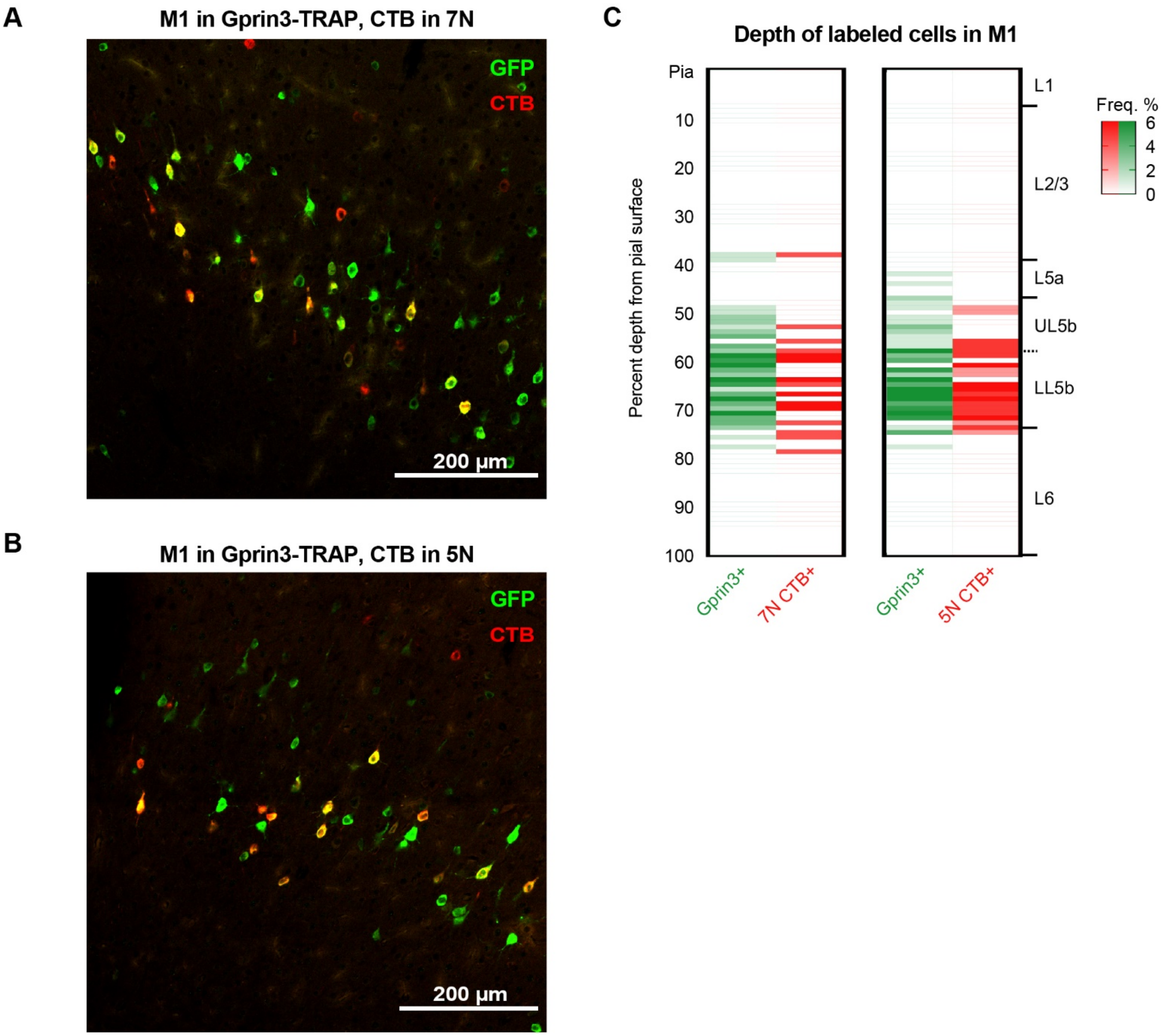


**Figure S3. A subset of Gprin3 cells show a projection to brainstem motor nuclei, related to Figure 2.** A) Immunostaining shows neuronal labeling in M1 of a Gprin3-TRAP mouse following a CTB injection in brainstem motor nucleus 7N. Green cells are GFP+ TRAP neurons, red cells are CTB+ 7N-projecting neurons. B) Immunostaining shows neuronal labeling in M1 of a Gprin3-TRAP mouse following a CTB injection in brainstem motor nucleus 5N. Green cells are GFP+ TRAP neurons, red cells are CTB+ 5N-projecting neurons. C) Frequency distribution of labeled neurons in M1 of Gprin3-TRAP mice following CTB injections in 7N (left) and 5N (right). Values reported as percent of cells found at each depth relative to the pial surface.

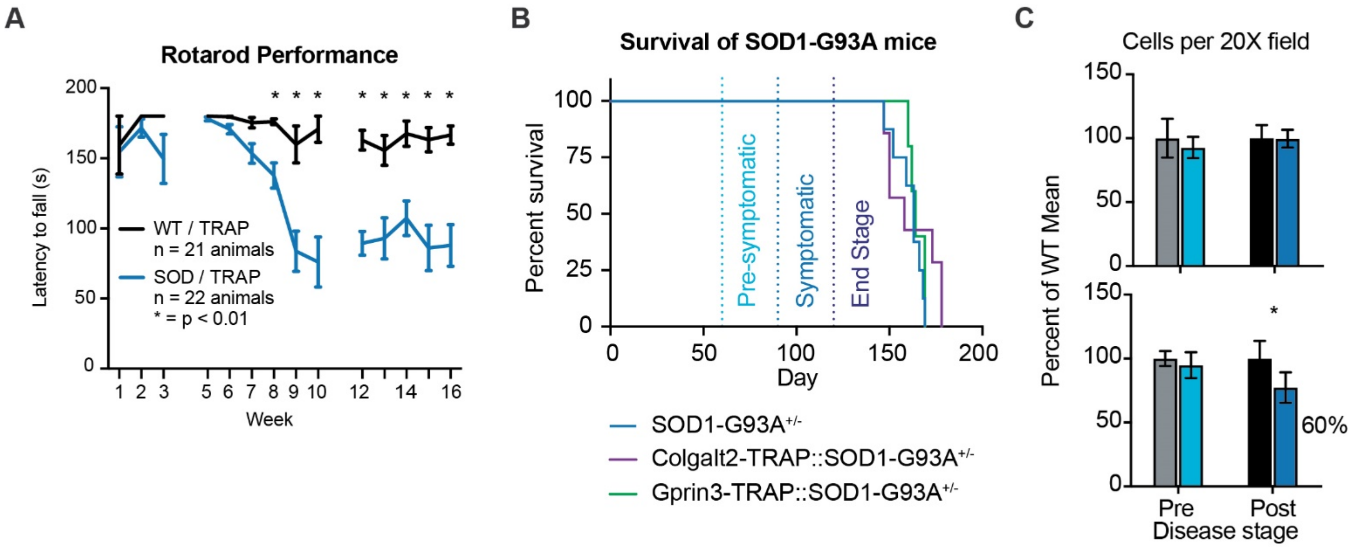


**Figure S4. SOD1-G93A::TRAP animals show standard symptom progression, related to Figure 3.** A) Survival curves for stock SOD1-G93A transgenic mouse line (grey, n = 8 animals), and Colgalt2-TRAP::SOD1-G93A (purple, n = 7 animals) and Gprin3-TRAP::SOD1-G93A (green, n = 5 animals) crosses. B) Latency to fall from rotarod across various stages of disease progression. Healthy WT animal performance (grey, n = 13 animals) and SOD1-G93A::TRAP animal performance (SOD; blue, n = 15 animals). Data represented as mean±SEM, * p < 0.0001 by unpaired t-test with Benjamini-Hochberg correction for multiple comparisons. C) Relative number of GFP+ cells across disease timepoints, average across all anterior-posterior coordinates of M1. Values are normalized to WT mean for the corresponding timepoint. * p-value < 0.05 by two-tailed unpaired t-test.

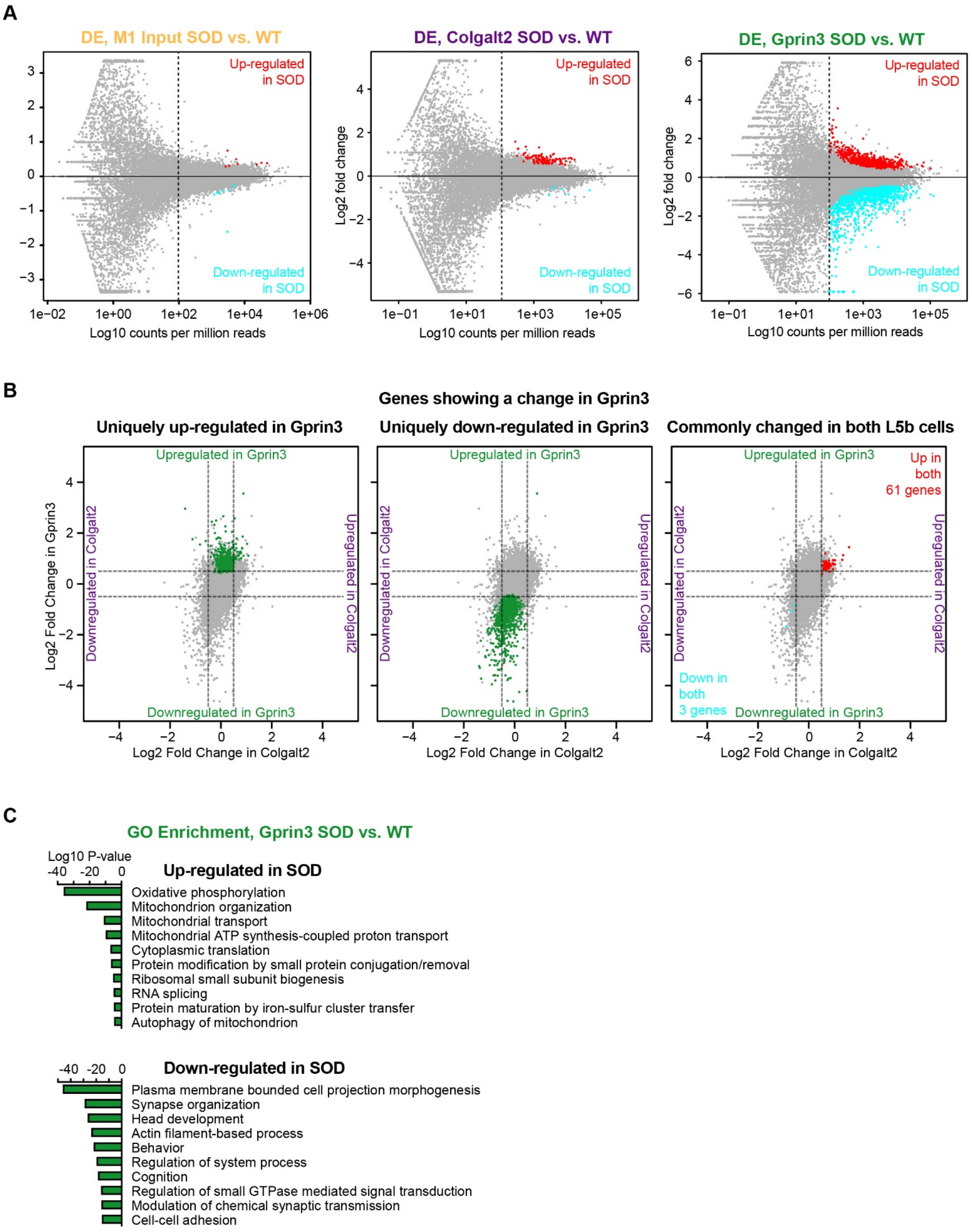


(**Figure S5** legend on next page)

**Figure S5. Differential expression analysis from SOD1-G93A::TRAP sequencing reveals cell type-specific changes in gene expression, related to Figures 4 and 5.** A) MA plots show differential expression between healthy WT and disease SOD sequencing samples for M1 input (left), Colgalt2 cells (center), or Gprin3 cells (right). Genes that showed significant downregulation in SOD are shown in cyan, and genes that were significantly upregulated in SOD are highlighted in red. B) Scatterplot shown in Fig. 4B, highlighting genes that were selectively up-regulated in Gprin3 cells (left), selectively down-regulated in Gprin3 cells (center), or regulated in both cell types (right). C) Bar graph shows more significantly enriched GO categories from GO analysis of Gprin3 SOD vs. WT DE. Values are plotted as log10 adjusted p-value for each category.

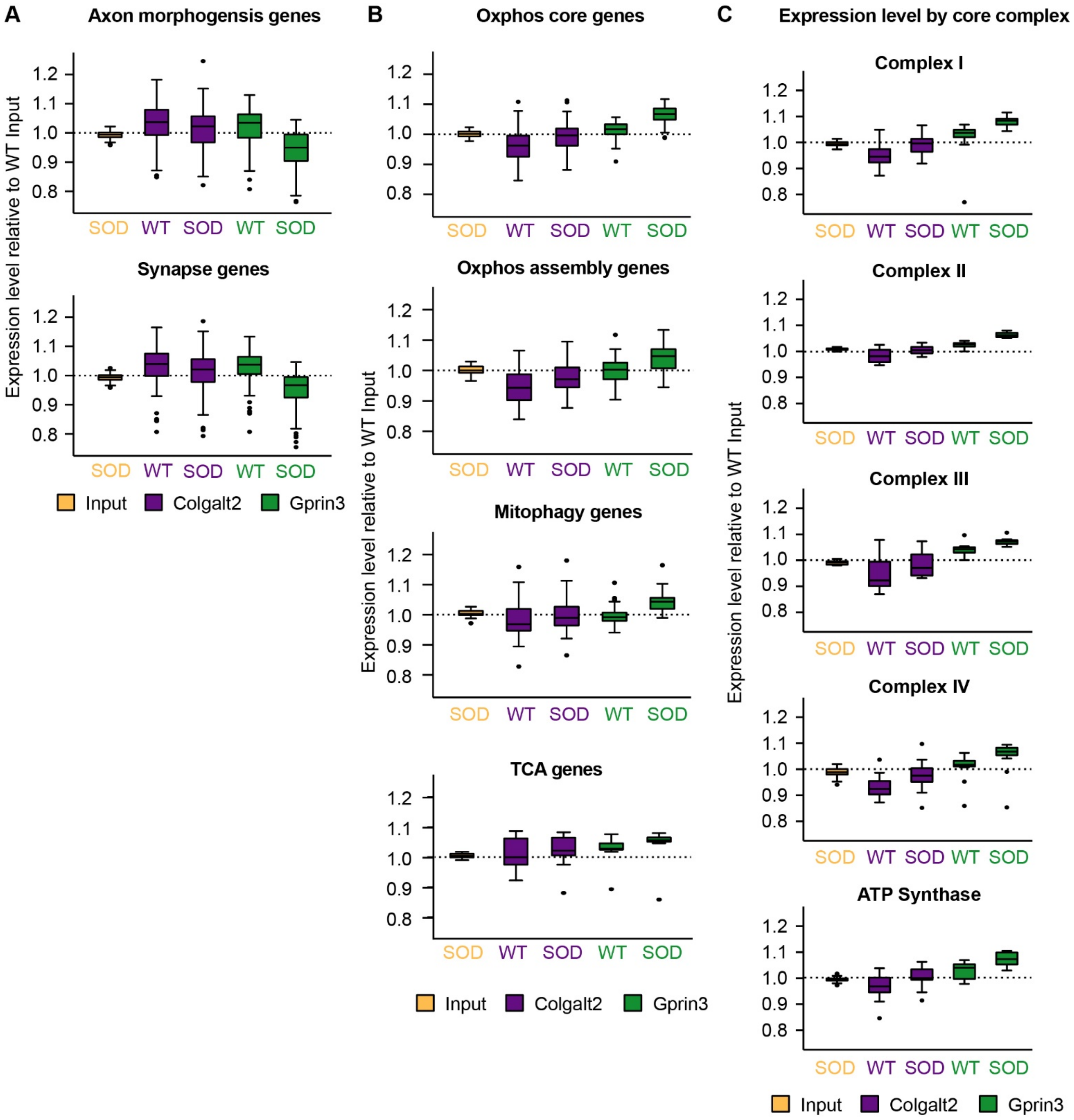


**Figure S6. Cell type-specific expression changes for functional groups of genes in SOD1-G93A, related to Figures 5 and 6.** A) SOD and WT CPM expression values normalized to WT M1 input CPM values for axon and synapse function and morphogenesis genes in M1 input, Colgalt2, and Gprin3 samples. B) SOD and WT CPM expression values normalized to WT M1 input CPM values for all genes that comprise the Oxphos core subunits, Oxphos assembly, mitophagy, and tri-carboxylic acid cycle (TCA) genes, in M1 input, Colgalt2, and Gprin3 samples. C) SOD and WT CPM expression values normalized to WT M1 input CPM values for genes that code for subunits of each complex of the electron transport chain, in M1 input, Colgalt2, and Gprin3 samples.

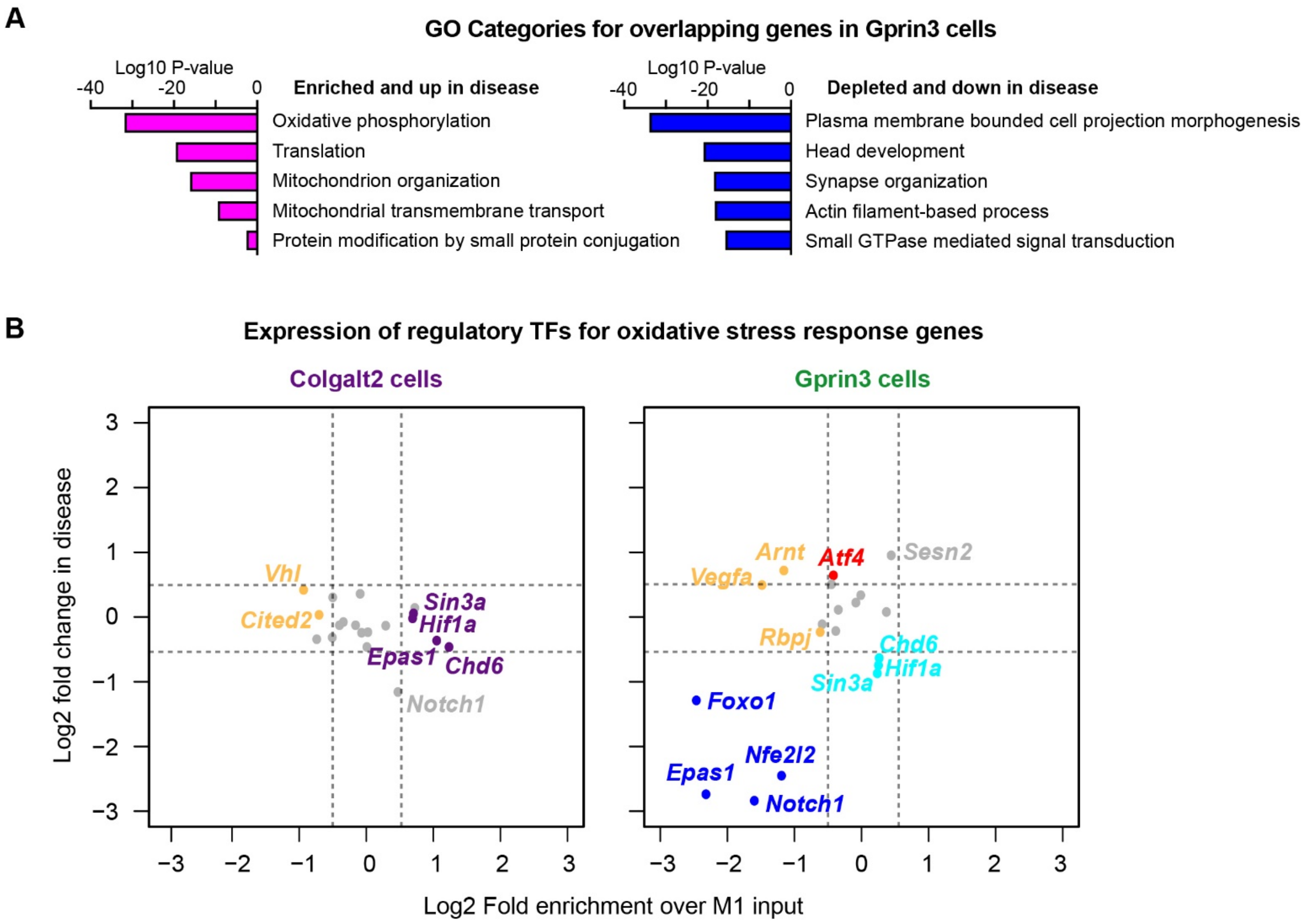


**Figure S7. In SOD1-G93A, Gprin3 cells modulate a set of genes that are already differentially enriched at baseline, related to Figure 7.** A) GO category enrichment for genes that showed enrichment in Gprin3 cells and also showed an upregulation in SOD1-G93A (magenta), and GO enrichment for genes that showed depletion in Gprin3 cells (aka. enrichment in Colgalt2 cells) and also showed a downregulation in SOD1-G93A (blue). Values shown are adjusted p-values. B) Scatterplots shows log2 fold enrichment from Colgalt2 vs. M1 input baseline DE (x-axis, left) and Gprin3 vs. M1 input baseline DE (x-axis, right) and log2 fold change from Gprin3 SOD vs. WT DE (y-axis, left and right) for transcription factors associated with activation of hypoxia and oxidative stress response pathways. Genes significantly depleted in Gprin3 or Colgalt2 samples relative to M1 input and significantly down-regulated in disease are highlighted in blue, genes down-regulated in disease but not enriched in either cell type are highlighted in cyan, genes enriched in Colgalt2 relative to M1 input but not changed in disease are highlighted in purple, genes that were enriched in M1 input over Colgalt2 or Gprin3 cells but were not changed in disease are highlighted in orange, and genes that were up-regulated in disease but not enriched in either cell type are highlighted in red.
